# The BTB transcription factors ZBTB11 and ZFP131 maintain pluripotency by pausing POL II at pro-differentiation genes

**DOI:** 10.1101/2020.11.23.391771

**Authors:** Görkem Garipler, Congyi Lu, Alexis Morrissey, Lorena S. Lopez-Zepeda, Simon E. Vidal, Begüm Aydin, Matthias Stadtfeld, Uwe Ohler, Shaun Mahony, Neville E. Sanjana, Esteban O. Mazzoni

**Author notes:** Equal contribution (Görkem Garipler, Congyi Lu). Corresponding authors: Neville E. Sanjana & Esteban O. Mazzoni.

## Abstract

In pluripotent cells, a delicate activation-repression balance maintains pro-differentiation genes ready for rapid activation. The identity of transcription factors (TFs) that specifically repress pro-differentiation genes remains obscure. By targeting ~1,700 TFs with CRISPR loss-of-function screen, we found that ZBTB11 and ZFP131 are required for embryonic stem cell (ESC) pluripotency. ZBTB11 and ZFP131 maintain promoter-proximally paused Polymerase II at pro-differentiation genes in ESCs. ZBTB11 or ZFP131 loss leads to NELF pausing factor release, an increase in H3K4me3, and transcriptional upregulation of genes associated with all three germ layers. Together, our results suggest that ZBTB11 and ZFP131 maintain pluripotency by preventing premature expression of pro-differentiation genes and present a generalizable framework to maintain cellular potency.

**One-sentence summary:** A Transcription Factor-wide CRISPR screen identifies ZBTB11 and ZFP131 maintaining pluripotency by pausing POL II at pro-differentiation genes

## Introduction

Early developmental progenitors and stem cells can maintain their fate while being able to differentiate rapidly. Pluripotency is an example of a transient state during early development, that can be recapitulated *in vitro* with ESCs isolated from the inner cell mass of blastocyst-stage embryos (Evans and Kaufman, 1981; Ying *et al.*, 2008). ESCs express pluripotency genes while keeping developmentally regulated genes repressed but ready for activation (Bernstein *et al.*, 2006; Mikkelsen *et al.*, 2007; Efroni *et al.*, 2008). The ability to rapidly activate pro-differentiation genes is critical for pluripotent cells to retain potency for lineage commitment yet differentiate rapidly during development. Although essential for understanding pluripotency and differentiation, dedicated TFs that guide repression mechanisms to specific pro-differentiation genes are still under investigation.

Controlling chromatin states and RNA Polymerase II (POL II) elongation dynamics are two major mechanisms to maintain plasticity for pro-differentiation gene expression. Bivalent genes contain nucleosomes modified by protein complexes associated with active (Trithorax group, TrxG) and repressive (Polycomb group, PcG) transcriptional states (Azuara *et al.*, 2006; Bernstein *et al.*, 2006; Vastenhouw and Schier, 2012). Similarly, reducing other chromatin modifiers such as histone deacetylase 1/2 (HDAC1/2) complexes leads to the upregulation of pro-differentiation genes (Kawamura *et al.*, 2005; Karantzali *et al.*, 2008; Dovey, Foster and Cowley, 2010; Jamaladdin *et al.*, 2014). Regulating POL II elongation dynamics by promoter-proximal pausing has emerged as a possible mechanism to repress pro-differentiation genes while preserving the ability to produce fulllength transcripts (Stock *et al.*, 2007; Ferrai *et al.*, 2017). Genes with promoter-proximal paused POL II can be rapidly transcribed, an ideal state for cell fate transitions during early development. In pluripotent stem cells, POL II pausing restricts the expression of pro-differentiation genes (Stock *et al.*, 2007; Adelman and Lis, 2012; Marks *et al.*, 2012). However, chromatin modifiers and POL II lack the sequence-specificity and must be recruited to a specific set of pro-differentiation genes that must be kept ready for activation.

Sequence-specific TFs are excellent candidates for recruiting and modulating chromatin complexes and POL II to regulate pro-differentiation genes. The pluripotency gene regulatory network (PGRN) relies on a set of interconnected TFs that engage in a positive feedforward loop to activate pluripotency genes (Li and Izpisua Belmonte, 2018). Moreover, the PGRN core TFs, OCT4, NANOG, and SOX2, directly bind to pluripotency and differentiation genes which must be actively maintained in a repressed state (Boyer *et al.*, 2005). *Ad hoc* repressive TFs that establish and control poised chromatin states in embryonic stem cells were postulated more than a decade ago (Boyer *et al.*, 2005; Creyghton *et al.*, 2008; Surface, Thornton and Boyer, 2010). However, a full understanding of sequence-specific repressive TFs that regulate the wide variety of genes required for the differentiation into all germ layers had remained elusive.

To rapidly dissect the gene regulatory network of ESCs, we developed a loss-of-function screen with a pooled CRISPR/Cas9 library targeting all annotated TFs in the mouse genome. The screen recovered known PGRN members (OCT4, NANOG, etc.) as well as two non-redundant ZBTB family TFs: ZBTB11 and ZFP131. ZBTB11 and ZFP131 co-occupy transcription start sites (TSS) of pro-differentiation genes along with TrxG, POL II, and NELF. Mutations of *Zbtb11* or *Zfp131* caused a decrease in NELF and an increase in H3K4me3 signal at their binding sites, which coincides with increased transcription of associated genes. As a result, ZBTB11 or ZFP131 loss causes transcriptomic changes associated with ESC differentiation into all three germ layers. Thus, we propose that ZBTB11 and ZFP131 play non-redundant roles as pluripotency gatekeepers to maintain developmental regulators in a repressed state but ready to be activated by restricting POL II elongation.

## Results

### A CRISPR loss-of-function screen to identify transcription factors required for pluripotency

To functionally identify TFs with essential roles during differentiation or fate maintenance, we designed and constructed a rapid and efficient CRISPR-Cas9 loss-of-function screen targeting all 1,682 annotated TFs in the mouse genome (“CRISPR TF library”) (Table S1) (Ravasi *et al.*, 2010; Meier, Zhang and Sanjana, 2017). The CRISPR TF library contains 10 single guide RNAs (sgRNAs) per TF and 1000 non-targeting sgRNAs (~5% of all sgRNAs) to serve as negative controls. We selected sgRNAs with predicted high on-target activity (Doench *et al.*, 2016), minimal potential for off-target (Hsu *et al.*, 2013), and those that target 5’ exons with annotated functional domains to enrich for loss-of-function mutations (Shi *et al.*, 2015) (Figure S1A-C, Table S1).

Because of their rapid cell cycle, *bona fide* ESCs outcompete differentiating cells or cells with compromised self-renewal abilities, a demonstrated advantage in identifying pluripotency and self-renewal genes (Ivanova *et al.*, 2006). Thus, ESCs transduced with sgRNAs targeting PGRN TFs should differentiate and be outcompeted by ESCs transduced with non-targeting sgRNAs and those targeting TFs that do not affect ESC state. A pilot competition assay using a non-targeting sgRNA versus sgRNAs targeting *Oct4 (Pou5f1), Nanog*, and *Klf5* (Figure S2A) revealed that cells transduced with sgRNAs targeting PGRN TFs were outcompeted starting by 5 days after transduction and were almost fully depleted by day 12 (Figure S2B). Therefore, we measured CRISPR TF library sgRNA representation on day 5, day 8, and day 12 after ESC transduction (Figure 1A). As expected, the sgRNA library diversity progressively decreased, as mutant cells for essential and pluripotency TFs were outcompeted over time (Figure 1B) (Shannon, 1948). To select candidate TFs, we used three different methods to identify significantly depleted sgRNAs: an empirical false discovery rate (FDR) cutoff derived from the embedded 1000 non-targeting sgRNAs controls (Chen *et al.*, 2015; Patel *et al.*, 2017), a mixed linear model with random intercepts using normalized sgRNA counts, and Model-based Analysis of Genome-wide CRISPR/Cas9 Knockout (MAGeCK) method (Li *et al.*, 2014).

**Figure 1.**
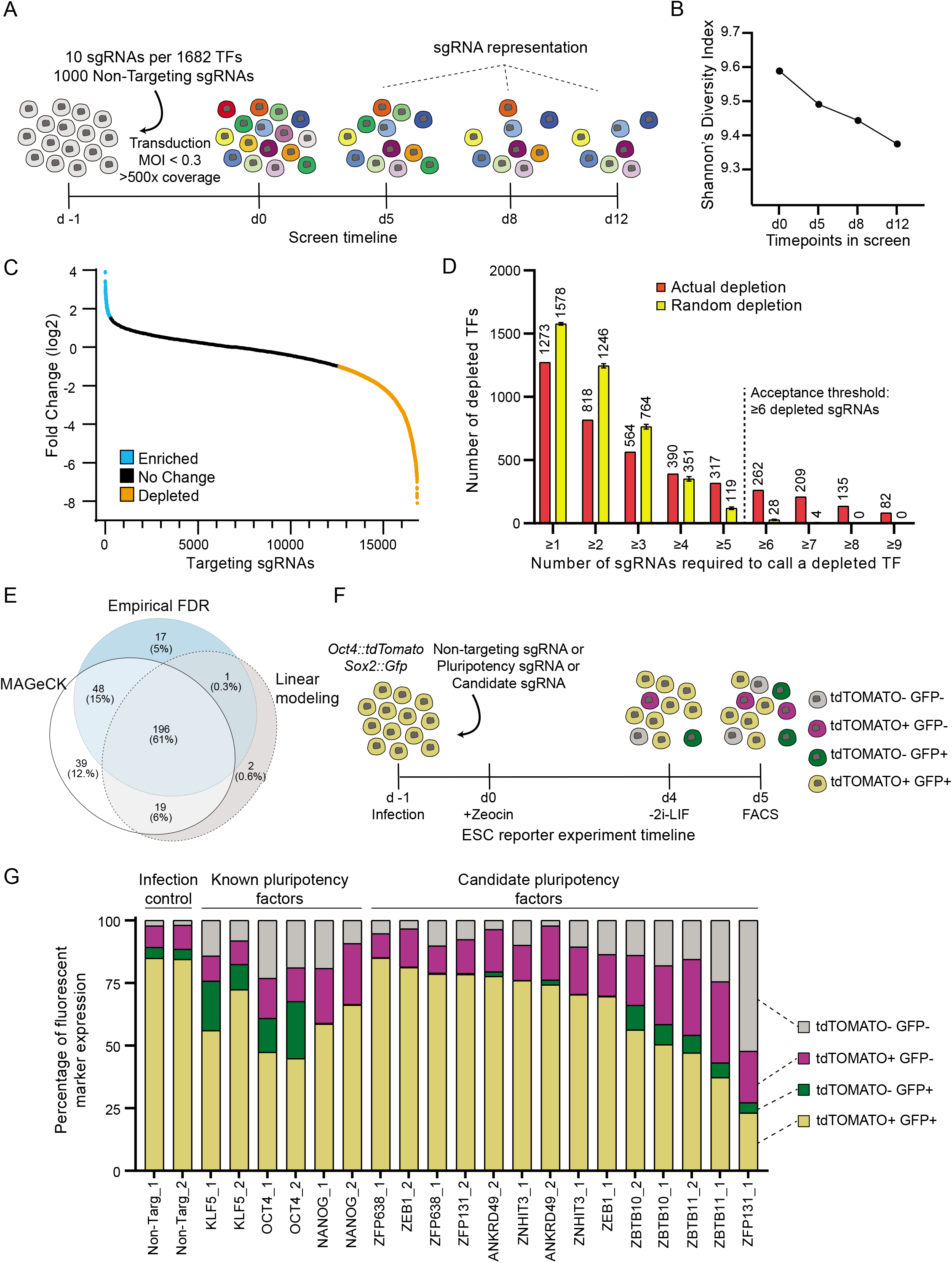
Identification of transcription factors required for pluripotency using the CRISPR TF library. **(A)** Transcription factor screen overview. Day −1: CRISPR TF library transduction to growing ESCs. Day 0: Zeocin selection begins. Day 5, 8, and 12: collection days and representation analysis. **(B)** Shannon’s diversity index (H) calculated based on the proportion of individual sgRNAs (pi) in the total number of sgRNAs in the pool (S) for each time point with sgRNA counts. The diversity score decreases as essential, and pluripotency targeting sgRNAs are lost from the population. **(C)** Log2 fold change of targeting sgRNAs on day 12 compared to the initial library representation. sgRNAs depleted more than the 50^th^ most depleted non-targeting sgRNA are in orange and enriched more than the 50^th^ most enriched non-targeting sgRNA are in blue. **(D)** The number of TF library sgRNAs depleted and mean sgRNAs number with random depletion calculated by 1000 permutation tests for the different number of sgRNA per gene. TFs with more than or equal to 6 depleted sgRNAs are unlikely to be false positives. **(E)** Venn diagram representation of the number of depleted TFs identified by three different methods: Log-fold change, linear modeling, and MAGeCK. 196 genes were picked as high confidence depleted TFs by all methods. **(F)** The workflow of secondary screen based on the *Oct4::tdTomato Sox2::Gfp* pluripotency reporter ESC line. Pluripotent ESCs are depicted as yellow while differentiating cells lose one or both reporter expression. **(G)** Percentage of cells expressing fluorescent reporters when transfected with a single sgRNA that is either non-targeting, targeting a known pluripotency TF, or candidate TF. 80% of cells infected with non-targeting sgRNA express both reporters. This number decreases in cells infected with sgRNAs targeting known pluripotency TFs and *Zbtb10, Zbtb11*, and *Zfp131*.

In the first approach, by measuring the depletion of the non-targeting sgRNAs, we set an empirical FDR cutoff of *q* < 0.05 and calculated the number of depleted sgRNAs per gene that exceeded this level (Figure 1C, S2C, D). Random permutation simulations demonstrated that at least 6 depleting or 3 enriching sgRNAs are required to confidently categorize a gene as a depleting or enriching, respectively (Figure 1D, S2E). In total, the empirical FDR method identified 262 depleting candidate TFs (Figure 1E). Although not the focus of this study, using an enrichment analysis, the empirical FDR method also identified 17 genes that increased self-renewal when mutated. These include the tumor suppressor *Trp53* (p53) and other genes which could provide growth advantage in cancers when mutated (Figure S3) (Cerami *et al.*, 2012; Gao *et al.*, 2013; Merkle *et al.*, 2017). In the second approach, the mixed linear model identified 218 depleting TFs (Figure 1E, S4A), and by using the degree of depletion placed candidate TFs within the PGRN hierarchy (Figure S4A, Table S2) (see methods). Finally, the MAGeCK approach identified 302 TF candidates as significantly depleted using robust-rank aggregation (Figure 1E, Table S2). In total, the three methods identified 196 TFs in common. As expected, this set included positive controls like essential genes and many known pluripotency factors. After removing these factors from this set, we selected 7 high-confidence novel candidate TFs (*Ankrd49, Zbtb10, Zbtb11*, *Zeb1, Zfp131, Zfp638, Znhit3)* with high scores and unknown functions to be tested for their putative role in pluripotency (Table S2).

### A secondary screen identifies ZBTB domain containing TFs as novel candidates for PGRN

Disrupting pluripotency or self-renewal decreases the ESCs’ proliferation rate. To discriminate between effects on pluripotency and self-renewal, we utilized a dual-color pluripotency reporter ESC line (Figure 1F). We generated an *Oct4:: tdTomato* and *Sox2::Gfp* pluripotency reporter line. When transduced with non-targeting sgRNAs, 85% of ESCs expressed both GFP and TDTOMATO (Figure 1G). In contrast, only 45 - 65% of cells transduced with sgRNAs targeting *Klf5, Nanog*, or *Oct4* expressed both fluorescent proteins under these culture and infection conditions (Figure 1G). We transduced the pluripotency dual-color reporter ESC line with two sgRNAs targeting each candidate TF (Figure 1F). To ensure robust, independent validation, we made sure that at least one sgRNA for each gene was not found in our original set of 10 sgRNAs per TF (Table S3). Transduction with sgRNAs targeting *Zbtb10, Zbtb11*, and *Zfp131* reduced OCT4 and SOX2 reporters similarly to the known pluripotency TFs (Figure 1G). In contrast, sgRNAs targeting *Zfp638, Ankrd49, Znhit3*, and *Zeb1* did not significantly reduce OCT4 and SOX2 reporter expression. We did not continue our investigation on these TFs since they are likely to control self-renewal or other ESC features, but not pluripotency.

A sgRNA competition assay between *Zbtb10, Zbtb11*, and Zfp131 targeting sgRNAs and non-targeting sgRNA (Figure S5A) demonstrated that *Zbtb11* and *Zfp131* sgRNAs depleted as fast as the core PRGN *Oct4* and *Nanog* sgRNAs (Figure S5B, S2A, B). *Zbtb10* depleted slower as *Klf5* (Figure S5B, S2A, B). The comparative depletion rate in the competition assay is in line with the original screen (Figure S4). Moreover, we performed a similar CRISPR TF screen using a library targeting all human TFs with 10 sgRNAs per gene and found that sgRNAs targeting the human homologs of *Zbtb10, Zbtb11*, and *Zfp131* are also depleted in human pluripotent stem cells (Figure S6). Thus, a rapid TF-wide screen combined in mouse and human ESC with a small-scale secondary screen using dual fluorescent reporter line was sufficient to identify novel candidate TFs for PGRN.

### Zbtb11 and Zfp131 are required for pluripotency

*Zbtb10*, *Zbtb11*, and *Zfp131* are C2H2 Zinc Finger BTB Domain TFs expressed in ESCs and during early mouse and human development (Boroviak *et al.*, 2018; Nowotschin *et al.*, 2019). To better characterize their role in pluripotency, we generated clonal ESC lines carrying null alleles for each TF. We introduced small frameshift deletions in the endogenous alleles while supplying an exogenous doxycycline-inducible HA-tagged rescue copy (Figure S5C). The addition of the HA-tag does not disturb ESC morphology or *Oct4* expression (Figure S5D, E). Doxycycline removal causes the extinction of the rescue construct, and consequently, ESCs only express null alleles for each candidate TFs (*Zbtb10Δ, Zbtb11Δ*, or *Zfp131Δ* thereafter). To validate the competitive disadvantage seen during the screen, we repeated the competition assay with H2B-GFP labeled wildtype (wt) ESCs plated in equal proportions with *Zbtb10Δ, Zbtb11Δ*, or *Zfp131Δ* ESCs (Figure S7A). Although there was a slight decrease, the H2B-GFP labeled wt ESCs did not significantly outcompete the *Zbtb10Δ* ESCs (Figure S7B). In contrast, the H2B-GFP labeled wt ESCs significantly outcompeted *Zbtb11Δ* and *Zfp131Δ* cells (Figure S7B).

Mouse ESCs grow in compact colonies and are typically cultured with LIF and with or without 2i compounds (MEK and GSK3 inhibitors) (Ying *et al.*, 2008). *Zbtb11Δ* and *Zfp131Δ* cells lost ESC morphology both with and without 2i (Figure 2A), and the number of OCT4 expressing cells decreased in *Zbtb11Δ* and *Zfp131Δ* (Figure S7C). On the other hand, *Zbtb10Δ* cells looked similar to wt cells (Figure 2A) and the number of OCT4-positive cells was mildly reduced in the *Zbtb10Δ* line (Figure S7C). Together, the characterization of clonal mutant lines confirms that ESCs require ZBTB11 and ZFP131, however, loss of ZBTB10 is not drastically affecting pluripotency features.

**Figure 2.**
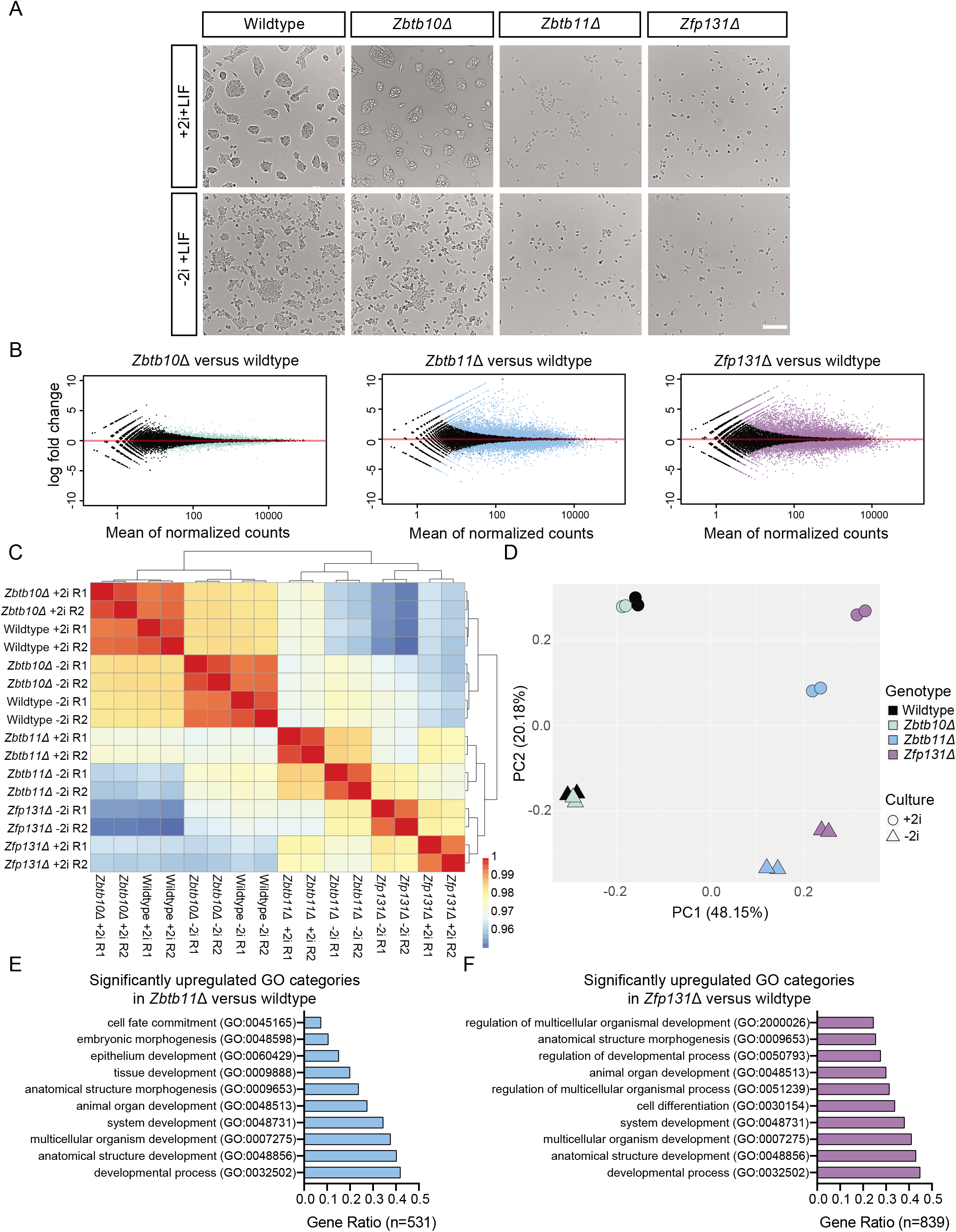
*Zbtb11Δ* and *Zfp131Δ* mutant cells express multiple lineage-specific markers. **(A)** *Zbtb10Δ, Zbtb11Δ* and *Zfp131Δ* colony morphology in +2i+LIF and −2i+LIF conditions 3d after doxycycline removal. *Zbtb11Δ* and *Zfp131Δ* cells lose typical ESC colony formation ability and adopt morphologies associated with differentiation. Scale bar = 200 μm. **(B)** Volcano plots of log2 fold change of transcripts expressed in *Zbtb10Δ, Zbtb11Δ*, and *Zfp131Δ* cells versus wildtype (wt) cells grown in the absence of doxycycline for 3d in −2i+LIF media (n=2). Significant changes (* p<0.05) marked in color for each genotype. *Zbtb11* and *Zfp131* mutations in ESCs induce aberrant gene expression. **(C)** Hierarchical clustering of transcriptomes of wildtype, *Zbtb10Δ, Zbtb11Δ*, and *Zfp131Δ* cells grown in the absence of doxycycline for 3d in −2i+LIF and +2i+LIF media. R1 and R2 denote two different replicates. *Zbtb11Δ* and *Zfp131Δ* cells have enough transcriptomic differences to cluster away from control and *Zbtb10Δ* cells in both culture conditions. **(D)** Principle component analysis of transcriptomes of wildtype, *Zbtb10Δ, Zbtb11Δ*, and *Zfp131Δ* cells grown in the absence of doxycycline for 3d in −2i+LIF and +2i+LIF media (n=2). Individual circles and triangles represent a single replicate. *Zbtb11* and *Zfp131* mutations induce profound transcriptomic changes separating mutant cells from wt and *Zbtb10* mutant cells along PC1. **(E)** GO-enrichment analysis of significantly upregulated genes in *Zbtb11Δ* (531) and **(F)** *Zfp131Δ* (839). *Zbtb11* and *Zfp131* mutations induce ESCs to express genes with a strong developmental signature.

### Loss of Zbtb11 and Zfp131 compromises pluripotent stem cell fate

Morphological changes of mutants described above led us to hypothesize that *Zbtb11Δ* and *Zfp131Δ* ESC would induce genes associated with cell differentiation. To test this hypothesis, we performed bulk RNA-seq on mutant cells from cells grown in LIF with and without and 2i. The transcriptome of *Zbtb10Δ* was similar to wt ESCs (Figure 2B, S8A). On the other hand, *Zbtb11* and *Zfp131* mutations induced severe transcriptome alterations in ESCs (Figure 2B, S8A). For example, *Zbtb11Δ* cells downregulated 301 and upregulated 795 genes, and *Zfp131Δ* cells downregulated 392 and 1189 upregulated genes without the 2i compounds.

Hierarchical clustering grouped each mutant according to their transcriptome (Figure 2C). *Zbtb10Δ* cells group together regardless of the culture conditions. On the other hand, *Zbtb11Δ* and *Zfp131Δ* separated from control ESCs independently of culture conditions (Figure 2C). Principal Component Analysis (PCA) of RNA-seq samples revealed a similar finding (Figure 2D). The two PCA dimensions explain 68.33% variance with the samples separated by genotype in the PC1 dimension (48.15% of the variance), while PC2 (20.18% of the variance) is the effect of removing 2i from the media. Interestingly, *Zbtb11* and *Zfp131* mutant cells are still sensitive to 2i removal, suggesting that they respond to Wnt and MEK signaling. Because of the slight growth disadvantage and the relatively small transcriptomic differences, we speculate that *Zbtb10* is not a central factor required for pluripotency. We conclude that *Zbtb11* and *Zfp131* are required for pluripotency because of the strong morphological and transcriptional changes in *Zbtb11Δ* and *Zfp131Δ* cells even in strong pluripotency maintaining conditions.

To understand ZBTB11 and ZFP131 in maintaining pluripotency, we characterized the set of dysregulated genes in their respective mutant lines. Notably, both mutant lines induced a strong differentiation signature in culture conditions with and without 2i compounds. Upregulated genes in *Zbtb11Δ* and *Zfp131Δ* cells are associated with cell differentiation GO-terms. Importantly, those include cell fate commitment, morphogenesis, and tissue development (Figure 2E, F, S8B-C). We also detected downregulated mitochondrial and other cellular biosynthetic and metabolic processes genes in *Zbtb11Δ* and *Zfp131Δ* ESCs (Figure S8D-F) (Wilson *et al.*, 2019). Thus, *Zbtb11* and *Zfp131* mutations caused profound transcriptional changes indicative of cell differentiation in the two most common ESC culture conditions.

### Zbtb11 and Zfp131 mutations induced ESCs to express genes associated with all three germ layers

We next asked if ZBTB11 and ZFP131 repress a unique differentiation trajectory each or if they repress the expression of genes associated with all three germ layers. Both mutants significantly downregulated pluripotency TFs and upregulated TFs associated with mesoderm, endoderm, ectoderm, and trophectoderm development compared to wt ESCs (Figure 3A). We did not detect any strong upregulation of genes associated with germline and primordial germ cells (Figure S8G). These results indicated that ZBTB11 and ZFP131 TFs are required for pluripotency maintenance with repressing pro-differentiation genes.

**Figure 3.**
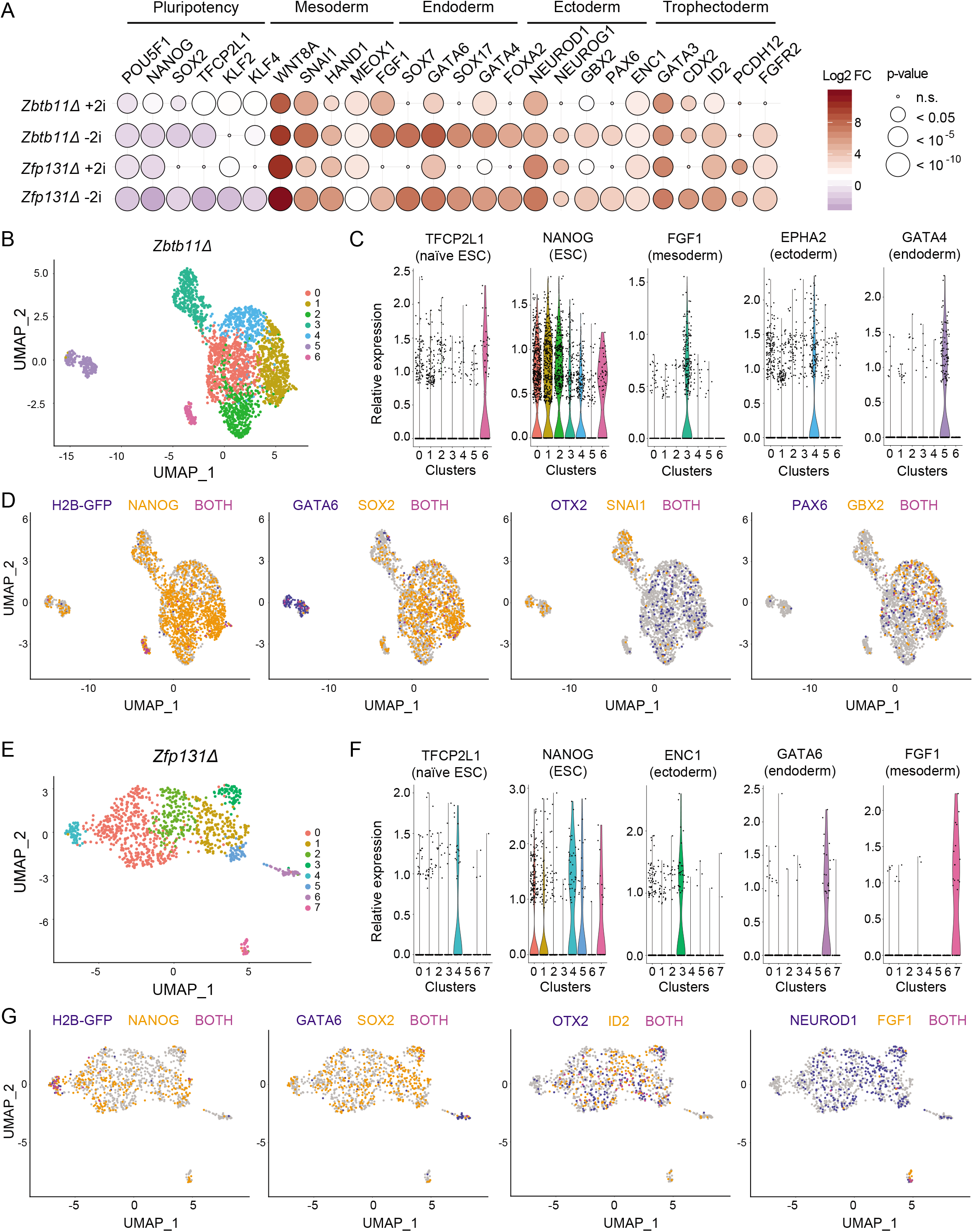
*Zbtb11Δ* and *Zfp131Δ* mutant cells differentiate into multiple lineages. **(A)** The log2 fold change of developmentally regulated TFs obtained from bulk RNAseq experiment in *Zbtb11Δ* and *Zfp131Δ* cells versus wt in +2i+LIF and −2+LIF conditions. (n.s. Not significant, * < 0.05, ** < 10^-5^, *** < 10^-10^) (n=2). *Zbtb11* and *Zfp131* mutations induce the expression of TFs associated with all three embryonic layers while downregulating pluripotency TFs. **(B)** Uniform manifold approximation and projection (UMAP) plot of *Zbtb11Δ* and *H2b::Gfp* (internal control) cells single-cell RNA-seq (n=1). **(C)** Violin plots of genes expressed in each cluster in *Zbtb11Δ* and *H2b::Gfp* cells. ESC, mesoderm, endoderm, ectoderm markers are expressed in distinct clusters. **(D)** Expression of genes associated with pluripotency and germ layers in *Zbtb11Δ* cells. *Zbtb11Δ* cells segregate in different clusters compared to *H2b::Gfp* and express developmentally regulated genes. **(E)** UMAP plot of *Zfp131Δ* and *H2b::Gfp* (internal control) cells single-cell RNA-seq (n=1). **(F)** Violin plots of genes expressed in each cluster in *Zfp131Δ* and *H2b::Gfp* cells. ESC, mesoderm, endoderm, ectoderm markers are expressed in distinct clusters. **(G)** Expression of genes associated with pluripotency and germ layers in *Zfp131Δ* and *H2b::Gfp* UMAP plots. Zfp131*Δ* cells segregate in different clusters compared to *H2b::Gfp* and express developmentally regulated genes.

The observed upregulation of developmental genes in bulk RNA-seq offered two alternative scenarios: mutant cells differentiate into multiple germ layer trajectories, or each cell simultaneously expresses genes of various germ layers. To discriminate between these two alternatives, we performed singlecell RNA sequencing (scRNA-seq). We mixed *Zbtb11Δ* or *Zfp131Δ* cells with H2B-GFP expressing wt ESCs to landmark control ESC expression in each experiment. H2B-GFP expressing wt naïve stem cells cluster away in both analyses with a high expression of stem cell markers (Cluster 6 in *Zbtb11Δ* Figure 3B and Cluster 4 in *Zfp131Δ* Figure 3E). *Zbtb11Δ* and *Zfp131Δ* each formed distinct clusters.

*Zbtb11Δ* cells were grouped in clusters with different germ layers markers (Figure 3C, D). For example, cluster 3 contained cells expressing ectoderm markers, while cluster 4 was associated with markers of mesoderm fate (Figure 3C, D). Cluster 5 included cells expressing genes related to endoderm (*Gata4*, *Gata6, Sox7). Zfp131Δ* clusters express genes associated with different germ layers with endoderm in cluster 6 and mesoderm markers in Cluster 7 (Figure 3F, G). *Zfp131Δ* cells upregulated ectodermal markers (*Otx2*, *Neurod1, Id2)* across several clusters (Figure 3G). Although *Zbtb11Δ* and *Zfp131Δ* clusters separated from ESCs, a portion of cells retained the expression of the pluripotency factors *Sox2* and *Nanog* (Figure 3D, G). This co-expression can be due to either capturing an early differentiation phase or aberrant expression. Together, neither *Zbtb11Δ* nor *Zfp131Δ* cells were grouped with wt ESCs nor formed a single cluster with each cell co-expressing genes from all germ layers. From the bulk and scRNA-seq analysis, we conclude that ZBTB11 and ZFP131 maintain the pluripotent state by actively repressing multiple developmental regulatory programs with each forming its own cluster and not one specific differentiation trajectory even with some aberrant gene combinations.

### ZBTB11 and ZFP131 bind to sites without classic repressive chromatin features

To gain insights into how ZBTB11 and ZFP131 maintain pluripotency and repress differentiation programs, we investigated the genomic binding of them in undifferentiated pluripotent cells. MultiGPS ChIP-seq analysis of HA-tagged exogenous copies identified 22,439 ZBTB11 sites and 3,635 ZFP131 binding sites (Mahony *et al.*, 2014). The majority of ZFP131 binding sites (78%) overlap with ZBTB11 (Figure 4A). BTB domains participate in homo and heterodimer formation, suggesting these two TFs could coregulate this set of genes (Stogios *et al.*, 2005). Both ZBTB11 and ZFP131 co-bound sites (Z11=Z131) and ZBTB11 only sites (Z11>Z131) contain a central motif similar to that bound by ETS family TFs: CCGGAA (Figure 4B). ZFP131 and ZBTB11 co-bound sites (Z11=Z131) and ZFP131 only sites (Z131>Z11) contain a central motif similar to that bound by bZIP TFs: TGACGTCA (Figure 4B). Using the GREAT algorithm (McLean *et al.*, 2010), we found that the majority of ZBTB11 and ZFP131 sites are proximal to TSSs. 46% of ZBTB11 binding and 72% of ZFP131 binding lies within 5 kb of nearest TSSs (Figure S8H, I). Comparing ZBTB11 and ZFP131 binding with ESC ATAC-seq experiments revealed that ZBTB11 and ZFP131 associate with accessible chromatin regions (Figure 4C) (Velasco *et al.*, 2017). These sites are also enriched in histone modifications associated with active transcription elements (H3K27ac) and promoter regions (H3K4me3) (Figure 4C). We noted these sites are also largely devoid of repressive histone modifications such as H3K9me3 and H3K27me3 (Figure 4C) (Bilodeau *et al.*, 2009). Therefore, ZBTB11 and ZFP131 bind at gene proximal sites containing an overall active chromatin signature.

**Figure 4.**
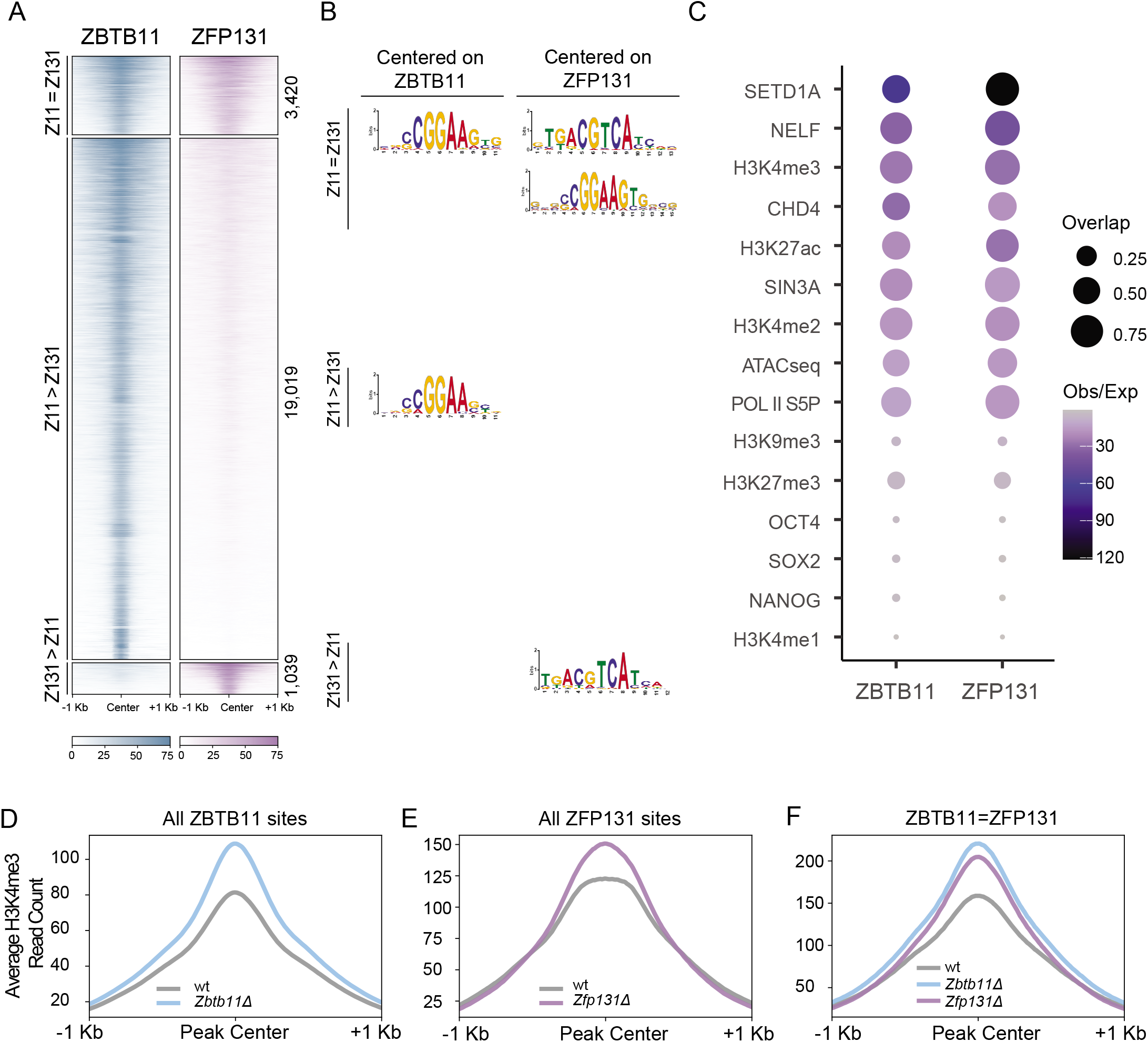
ZBTB11 and ZFP131 bound sites gain an H3K4me3 signal in *Zbtb11Δ* and *Zfp131Δ* mutant cells. **(A)** ChIP-seq heatmap for ZBTB11 and ZFP131 binding on the ESC genome (n=2). Categories were identified with ZBTB11 and ZFP131 cobinding (Z11=Z131), differentially enriched ZBTB11 binding (Z11>Z131) and differentially enriched ZFP131 binding (Z131>Z11). **(B)** Enriched motifs discovered via MEME-ChIP for each category of ZBTB11 and ZFP131 binding. Motif discovery was performed with centering to the peak at ZBTB11 and ZFP131 for three differential binding categories. **(C)** Percentage of overlap between ZBTB11 and ZFP131 with pluripotency factors, known regulatory complex members, histone modifications, and ATAC-seq signal. ZBTB11 and ZFP131 bind at sites with active chromatin states, accessible and enriched with TrxG, Pol II S5P, NELF and histone deacetylase members. ZBTB11 and ZFP131 do not co-bind with pluripotency TFs but bind to regions with active chromatin signatures. **(D)** Metagene plots for H3K4me3 difference at ZBTB11 bound sites in *Zbtb11Δ* versus wt (n=2), **(E)** at ZFP131 bound sites in *Zfp131Δ* versus wt (n=2), and **(F)** at ZBTB11 and ZBTB131 cobound sites in *Zbtb11Δ* and *Zfp131Δ* versus wt. *Zbtb11* and *Zfp131* mutations result in an H3K4me3 gain at ZBTB11 and ZFP131 binding sites.

We considered several hypotheses that could explain how *Zbtb11* and *Zfp131* maintain pluripotency. The ZBTB family TF ZBTB3 occupies the ESC genome with core pluripotency factors to modulate their activity (Ye *et al.*, 2018). However, low ZBTB11 and ZFP131 binding overlap (<5%) with OCT4, SOX2, and NANOG do not support the hypothesis of co-regulation between ZBTB11 or ZFP131 and pluripotency factors (Figure 4C). ZBTB TFs are also known to interact with the subunits of histone deacetylase complexes (David *et al.*, 1998; Maeda, 2016; Masuda *et al.*, 2016). Both ZBTB11 and ZFP131 overlap with sites occupied by CHD4 and SIN3A, known components of different histone deacetylase complexes (Figure 4C, Figure S9A-B) (Laherty *et al.*, 1997; Xue *et al.*, 1998). While there were small CHD4 and SIN3A decreases at ZBTB11 and ZFP131 sites in *Zbtb11Δ* or *Zfp131Δ* cells (Figure S9C-F), these areas lost H3K27ac, which is contrary to expectation when losing HDACs (Figure S9G, H). Overall, H3K27ac deposition did not correlate with significantly decreased or increased CHD4 and SIN3A in *Zbtb11Δ* or *Zfp131Δ.* Thus, these results do not support a direct role in HDAC recruitment.

### ZBTB11 and ZFP131 prevent TrxG activation

ZBTB11 and ZFP131 TSS-proximal binding suggest that they could maintain a poised state at pro-differentiation genes. However, ZBTB11 or ZFP131 sites do not overlap with high levels of H3K27me3, indicating that their target genes do not rely on PRC2 repression (Figure 4C). Therefore, our data do not support ZBTB11 and ZFP131 binding as a mechanism to recruit PcG to maintain the bivalent state of developmentally regulated genes in ESCs.

ZBTB11 and ZFP131 binding overlaps with H3K4me3, which is deposited by TrxG (Figure 4C). Among the TrxG H3K4me-depositing enzymes, SETD1A is expressed and required at early developmental states until inner cell mass formation in development (Bledau *et al.*, 2014). SETD1A binding overlaps significantly with ZBTB11 and ZFP131 in wt ESCs (Figure 4C, 5A, 5B) (Sze *et al.*, 2017). Removing ZBTB11 or ZFP131 induced a significant increase of H3K4me3 at their binding sites (Figure 4D, E). This increase was even more prominent at sites where ZBTB11 and ZFP131 binding overlaps (ZBTB11=ZFP131) (Figure 4F). Importantly, most of the regions that gained H3K4me3 in *Zbtb11Δ* and *Zfp131Δ* cells already contained SETD1A binding in the wt state (Figure 5C-E). Genes that were bound by ZBTB11 or ZFP131 that gained H3K4me3 and mRNA upregulation in mutant lines are strongly associated with differentiation GO terms (Figure 5F, G). These genes included *Gata6, Sox7, Id3, Snai1, Sox17, Onecut1, Nkx2.2*, and *Gata3* pro-differentiation TFs. Therefore, our results suggest that ZBTB11 and ZFP131 bind to a subset of pro-differentiation genes that already contain SETD1A the TrxG complex in ESCs. Upon ZBTB11 or ZFP131 removal, these sites gained H3K4me3 with the associated gene transcription, particularly at ZBTB11 and ZFP131 co-bound sites. We conclude that ZBTB11 and ZFP131 are the sequence-specific TFs that maintain sufficient repression force to prevent premature pro-differentiation gene upregulation and spontaneous pluripotent stem cell differentiation.

**Figure 5.**
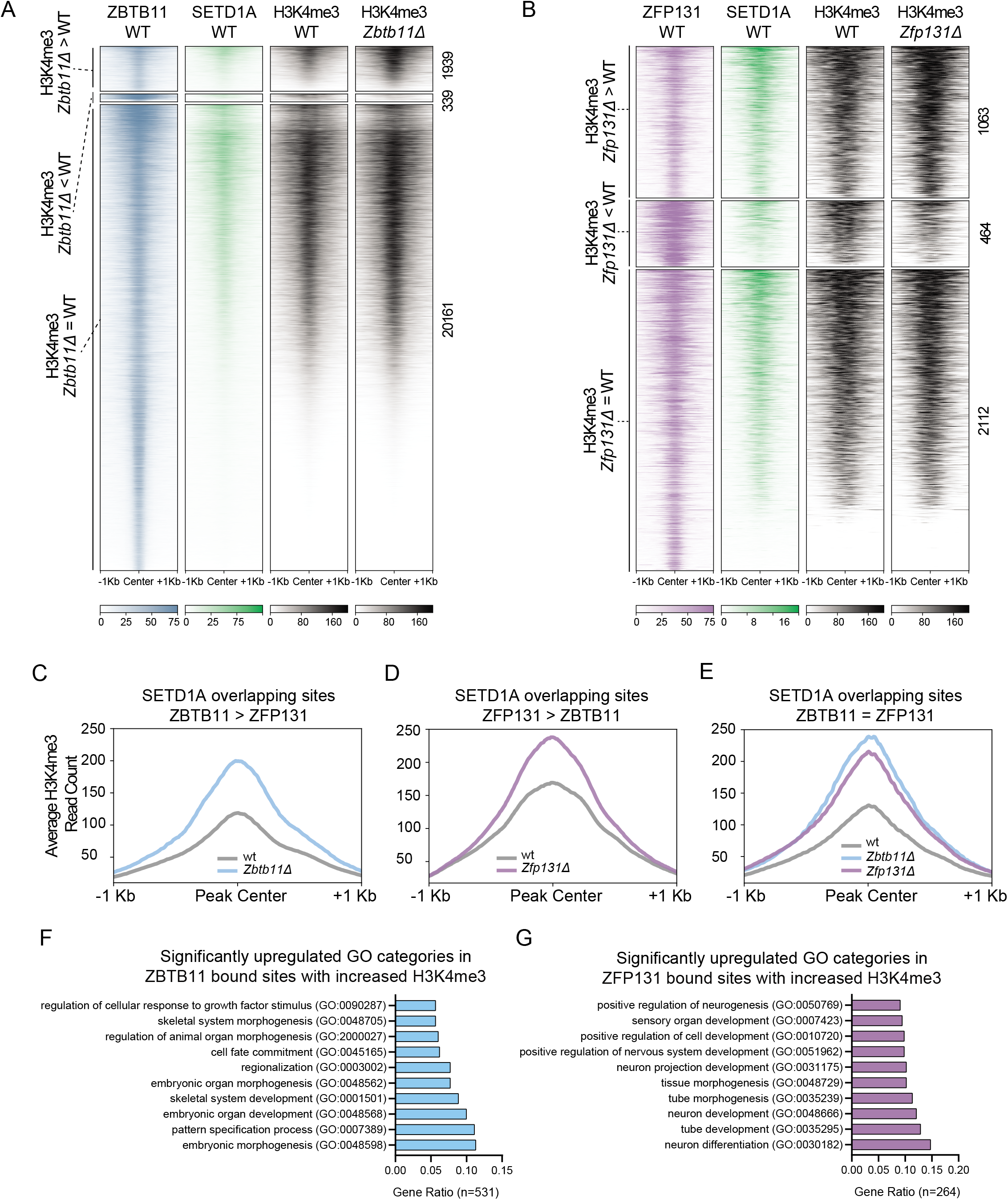
SETD1A occupancy at ZBTB11 and ZFP131 sites. **(A)** ChIP-seq heatmap for ZBTB11 and **(B)** ZFP131 overlap with SETD1A and H3K4me3. Categories are determined according to gain, loss or no change in H3K4me3 signal between mutant and wt cells (n=2 for all). ZBTB11 and ZFP131 coocupy ESC genome with SETD1A. **(C)** Metagene plots for H3K4me3 signal difference in ZBTB11 and SETD1A co-bound sites in *Zbtb11Δ* versus wt, **(D)** in ZFP131 and SETD1A co-bound sites in *Zfp131Δ* versus wt, **(E)** in ZBTB11, ZFP131 and SETD1A co-bound sites in all three lines (n=2 for all). ZBTB11 and ZFP131 removal induced H3K4me3 gain at SETD1A present sites in wt cells. **(F)** GO-enrichment analysis of significantly upregulated genes with ZBTB11 binding and H3K4me3 increase (n=531) and **(G)** with ZFP131 binding and H3K4me3 increase (n=264). Gene Ratio is calculated with the number of occurrences in specific GO categories divided by the total amount of genes. GO-terms are associated with differentiation.

### ZBTB11 and ZFP131 regulate promoter-proximal POL II pausing at pro-differentiation genes

Low H3K27me3 signal at ZBTB11 and ZFP131 bound sites indicate that these TFs are not regulating bivalent chromatin, therefore we sought to investigate if ZBTB11 and ZFP131 regulate POL II elongation dynamics. POL II carboxyl-terminal domain (CTD) contains repeats of the Y_1_S_2_P_3_T_4_S_5_P_6_S_7_ amino acid sequence, which is unphosphorylated during POL II recruitment to the TSSs (Hsin and Manley, 2012). POL II is loaded onto TSSs, forming a complex with the negative elongation factor (NELF) among other factors, pausing proximal to the TSS (Yamaguchi *et al.*, 1999; Muse *et al.*, 2007; Zeitlinger *et al.*, 2007). Following initial elongation, POL II is phosphorylated at serine 5 (S5P), but further elongation is enabled by serine 2 phosphorylation (S2P) and with NELF removal (Egloff, Dienstbier and Murphy, 2012).

Both ZBTB11 and ZFP131 overlap with serine 5 phosphorylated POL II (POL II S5P) (Figure 4C, 6A). Moreover, these sites also contain NELF (Figure 4C, 6A). ZBTB11 and ZPF131 removal decreased NELF binding at these sites (Figure 6B). Genes containing POL II S5P, NELF, and either ZBTB11 or ZFP131 were upregulated in *Zbtb11Δ* and *Zfp131Δ* cells (Figure 6C-D). In agreement with chromatin phenotypes, the transcription of genes associated with co-bound sites (ZBTB11=ZFP131) was strongly upregulated (Figure 6E-F). These results suggest that ZBTB11 or ZFP131 removal causes NELF displacement, allowing POL II to engage in transcription elongation, leading to transcriptional upregulation.

**Figure 6.**
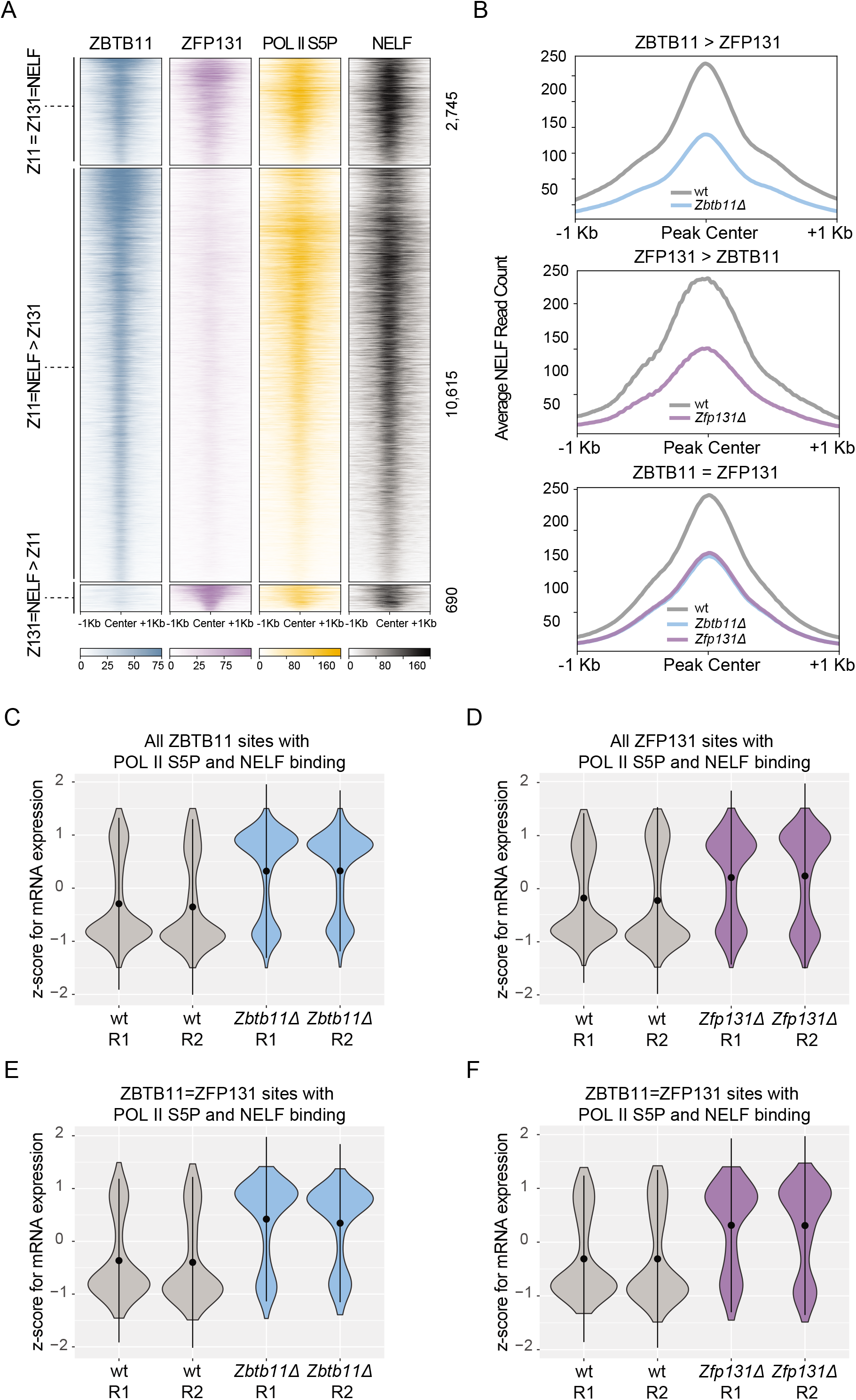
POL II S5P and NELF promoter-proximal pausing complex overlaps ZBTB11 and ZFP131 sites. **(A)** ChIP-seq heatmap for ZBTB11 and ZFP131 overlap with POL2 S5P and NELF. Categories are determined according to ZBTB11 only, ZFP131 only and cobound regions (n=2 for all). ZBTB11 and ZFP131 bind at sites with POL II S5P and NELF in ESCs. **(B)** Metagene plots for NELF signal difference in ZBTB11 only, ZFP131 only and cobound regions (n=2 for all). ZBTB11 and ZFP131 sites lose NELF in *Zbtb11* and *Zfp131* mutant ESCs. **(C)** Violin plots for z-score mRNA expression of genes that are ZBTB11 bound along with NELF and POL II S5P with H3K4me3 increase in *Zbtb11* mutant (n=1191), **(D)** ZFP131 bound along with NELF and POL II S5P with H3K4me3 increase in *Zfp131* mutant (n=915), **(E)** ZBTB11 and ZFP131 bound along with NELF and POL II S5P with H3K4me3 increase in *Zbtb11* mutant (n=216), and **(F)** ZBTB11 and ZFP131 bound along with NELF and POL II S5P with H3K4me3 increase in *Zfp131* mutant (n=216). ZBTB11 and ZFP131 genes with paused POL II become transcribed in *Zbtb11* and *Zfp131* mutant ESCs.

Together, our results show that ZBTB11 and ZFP131 bind proximally to developmentally regulated genes to regulate POL II elongation dynamics. These genes contain SETD1A, low H3K4me3 levels, and promoter-proximal paused POL II along with NELF (Figure 7) but have restricted expression in ESCs. Upon ZBTB11 and ZFP131 removal, their target genes gain H3K4me3, decommission NELF, and upregulate transcription (Figure 7). ZBTB11 and ZFP131 are not redundant since individual mutants have phenotypes and associated genes at co-bound sites are also subjected to this regulation. Thus, ZBTB11 and ZFP131 provide repression with POL II pausing to maintain pro-differentiation genes at a delicate activation-repression balance to preserve pluripotency.

**Figure 7.**
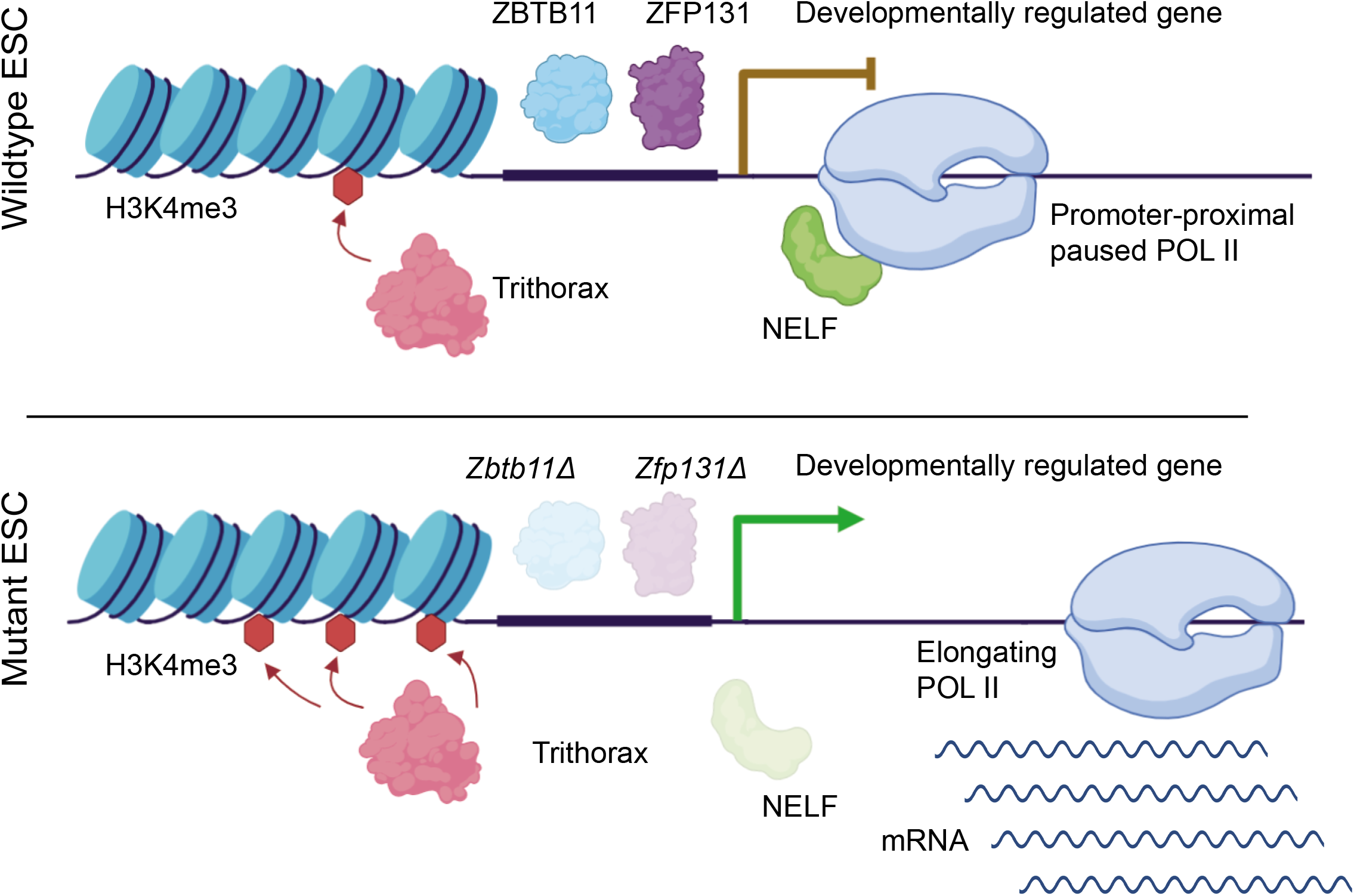
ZBTB11 and ZFP131 maintain developmentally regulated genes at a poised state. Schematic overview of the mode-of-action of ZBTB11 and ZFP131 in ESCs. ZBTB11 and ZFP131 bind proximal to developmentally regulated genes. These genes contain promoter-proximal paused POL II and NELF, and chromatin features associated with poised states. Upon ZBTB11 and ZFP131 removal, TrxG increases activity, and POL II transcribes pro-differentiation genes causing ESC to differentiate and exit their pluripotent state.

### Conclusions

Transcription factors are critical regulators of cell fate, typically playing instructive roles in cell fate maintenance and differentiation. Decades of screens and studies have yielded a profound understanding of the functions of many TFs. However, there is still an overwhelmingly large fraction of TFs with no described role, even in one of the most molecularly well-characterized cellular states: pluripotent stem cells. To gain insights into TF-driven gene regulatory networks, we designed a loss-of-function CRISPR-Cas9 screen targeting all annotated TFs in the genome and deployed this unique tool to identify key TFs that regulate pluripotency. We expect this strategy to shed light on other TF-dependent regulatory processes such as differentiation and development of diverse cell lineages.

The PGRN is composed of interlocked activating TFs. At its core, the OCT4-SOX2 complex binds with other partners, including NANOG, in a strong pluripotency positive feedback loop (Boyer *et al.*, 2005; Kim *et al.*, 2008; Festuccia *et al.*, 2012). OCT4 and SOX2 bind proximally to pro-differentiation genes (Boyer *et al.*, 2005; Jaenisch and Young, 2008); however, these TFs are not *ad hoc* repressors suggesting additional TFs with repressive functions are required. With the TF-focused loss-of-function screen, we uncovered a novel role for the BTB-domain ZBTB11 and ZFP131 TFs. Pluripotency requires *Zbtb11* and *Zfp131*, and their mutations upregulate the expression of genes associated with differentiation to all three germ layers. ZBTB11 and ZFP131 do not cooccupy ESC chromatin with core pluripotency TFs, thus unlikely to form a repressive complex with OCT4-SOX2. ZBTB TFs generally have repressive roles with BTB domains interacting with HDACs, nuclear co-repressors (*NCoR1*), silencing mediator of retinoic acid and thyroid hormone receptor (*NCoR2*/SMRT), and *Sin3A* (Bardwell and Treisman, 1994; Albagli *et al.*, 1995; Melnick *et al.*, 2002; Ahmad *et al.*, 2003; Ghetu *et al.*, 2008). However, ZBTB11 and ZFP131 binding neither overlap with NCOR nor impact histone deacetylase activity. Finally, ZBTB11 and ZFP131 binding does not overlap with bivalent genes or PcG deposited H3K27me3 signal, suggesting they have no role in bivalent chromatin state regulation.

Rapid gene activation during cell differentiation can be controlled by regulating POL II pausing and elongation (Guenther *et al.*, 2007; Marks *et al.*, 2012). Characterizing POL II pausing in wt and POL II pausing-deficient cells suggested that around 40% of all developmental regulators contain paused POL II at their TSSs (Williams *et al.*, 2015). We propose that ZBTB11 and ZFP131 prevent aberrant induction of pro-differentiation genes in pluripotent stem cells by establishing or maintaining POL II pausing (Figure 7). Genetic ablation of NELF does not cause a differentiation phenotype, whereas resulting in profound effects in cell growth, metabolism, and kinase activity (Williams *et al.*, 2015). We speculate that these phenotypic differences are due to ZBTB11 or ZPF131 affecting a subset of NELF bound genes, contrary to the pleiotropic effect seen in the NELF mutant. Another ZBTB TF family member, ZBTB17 (MIZ-1) was previously described to repress a set of *Hox* genes in pluripotent stem cells (Varlakhanova *et al.*, 2011). It would be interesting to test if ZBTB17 uses a similar repression mechanism as ZBTB11 and ZFP131.

The BTB domain can form homo and heterodimers (Stogios *et al.*, 2005). The similar phenotype of both mutants and the binding overlap, particularly for ZFP131, suggest that they might coregulate a set of developmentally regulated genes. Moreover, a larger overlap is conceivable if a higher number of sites could be recovered from ZFP131 ChIP. The fact that both genes were recovered from the functional screen with a similar phenotype for each mutant suggests that these TFs perform similar but non-redundant functions to maintain pluripotency.

*Zbtb11* and *Zfp131* are expressed in pluripotent stem cells and at early pluripotent developmental stages (Boroviak *et al.*, 2018; Nowotschin *et al.*, 2019). Although pluripotency can be maintained indefinitely by chemical signals *in vitro*, it is a transient *in vivo* state. We speculate that ZBTB11 and ZFP131 safeguard the transient state of pluripotency by preventing premature activation of differentiation during development. Broadly, the temporal gene activation in pluripotent and multipotent states is tightly regulated during vertebrate development. Since only a fraction of all TFs expressed during development are functionally characterized, we predict that TFs similar to ZBTB11 and ZFP131 have equivalent roles in maintaining poised genes in transient multipotent progenitors at later developmental stages.

## Acknowledgments

This work was supported by the NICHD (R01HD079682) to E.O.M. N.E.S. is supported by New York University and New York Genome Center startup funds, National Institutes of Health (NIH)/National Human Genome Research Institute (grant nos. R00HG008171, DP2HG010099), NIH/National Cancer Institute (grant no. R01CA218668), Defense Advanced Research Projects Agency (grant no. D18AP00053), the Sidney Kimmel Foundation, the Melanoma Research Alliance, the Cancer Research Institute, and the Brain and Behavior Foundation. A.M. is supported by the CBIOS training grant from NIGMS (T32GM102057). The Mahony lab is supported by NIGMS (R01GM125722) to S.M. L.S.L.Z. is supported by the MDC-NYU exchange program. The Stadtfeld lab is supported by Tri-SCI 2019-026 from Tri-Institutional Stem Cell Initiate and R01GM121994 to M.S.

The authors would like to thank: the Mazzoni lab members, Claude Desplan, Danny Reinberg, and Ana Pombo for feedback and constructive comments; the NYU Genomics Core facility for technical support and data processing; Disi An for establishing the H2B::GFP reporter line; Silvia Velasco for generating input control for ChIP-seq in ESCs; Roland Schwarz for helpful input on the linear models; Albert Tan for quantifying competition experiments.

## Author Contributions

G.G performed the mouse CRISPR TF screen, secondary screen, established inducible knockout lines, performed RNA-seq, sc-RNA-seq, ChIP-seq experiments, competition assay, imaging experiments, and the analysis of RNA-seq and sc-RNAseq data. G.G and C.L cloned the human and mouse CRISPR TF libraries. C.L performed the human CRISPR TF screen. A.M performed the analysis of ChIP-seq data and set up the computational analysis framework for analysis and interpretation with the support from S.M. and G.G. L.S.L.Z and U.O. developed the time-course linear modeling for CRISPR perturbations. L.S.L.Z identified significant hits using linear modeling. S.E.V. generated *Oct4::tdTomato Sox2::Gfp* cell line with support from M.S. G.G and B.A. have established initial scRNAseq set up for ESCs. N.E.S designed the human and mouse CRISPR TF libraries. G.G, C.L., N.E.S, and E.O.M conceived the experiments. G.G and E.O.M co-wrote the original draft of the manuscript. All authors read and approved the final manuscript.

## Data Availability

All sequencing data (ChIP-seq, RNA-seq, scRNA-seq, GeCKO-seq) has been deposited at the GEO database under accession number GSE160966. We performed re-analysis of data sourced from GEO database entries GSE24165, GSE18371, GSE11724, and GSE11172.

## Methods

### Generation of stem cell lines and culture conditions

A17 E14Tg2a line was used for the initial CRISPR TF screen experiments (Iacovino *et al.*, 2011). *Zbtb11* and *Zfp131* were amplified from cDNA of ESCs using Zbtb11-fwd (5’ ATGTCAAGCGAGGAGAGCTACC), Zbtb11-rvs (5 CTCTGCCTCTGGCATATGTGC), Zfp131-fwd (5’ ATGGAGGCTGAAGAGACGATGG) and Zfp131-rvs (5’ TTCTAAAACCGGCAGAGATGTCC) primers. The cDNA of *Zbtb10* was amplified from Origene (MR214195L4) *Zbtb10* mouse tagged ORF clone using Zbtb10-fwd (5’ ATGTCGTTCAGTGAGATGAACCG) and Zbtb10-rvs primer (5’ TCCAGAGACATAAACGCCTCC). C-terminus of *Zbtb10, Zbtb11*, and *Zfp131* cDNAs were fused to 1x hemagglutinin (HA) tag and cloned into p2lox plasmid using In-fusion (Clontech) cloning. To avoid sgRNAs from both landing on exogenous alleles and Cas9 from cutting at the PAM site, the sgRNA docking site on the BTB domain sequence of *Zbtb10, Zbtb11* and *Zfp131* was altered without changing the amino acid sequence (Figure S6A). Resulting CRISPR-proof alleles in p2lox-*Zbtb10*-HA, p2lox-*Zbtb11*-HA, p2lox-*Zfp131*-HA were nucleofected into mouse ESCs grown in 1 μgml^-1^ doxycycline (Sigma, D9891) to induce iCre expression for 16h. Successfully inserted transgenes were selected using G418 selection (400 μg ml^-1^, Cellgro) and characterized for expression of tagged transgenic proteins with HA staining (anti-HA, ab9110).

To knock out endogenous alleles of *Zbtb10, Zbtb11*, and *Zfp131*, inducible *Zbtb10-HA, Zbtb11-HA* and *Zfp131-HA* lines were nucleofected with CMV-GFP and pTF vector (Figure S1D). Two pTF vectors with sgRNAs targeting 100 bp apart from each other with *Zbtb10* sgRNAs (5’ TCTTTGTGATGTCAGCATTG and 5’ AGAAACGGCTGCCTGCAACC), *Zbtb11* sgRNAs (5’ AGCGCACAAGTCTGTCCTCT and 5’ AGGAGCAGTTTCTAGTCACG), and *Zfp131* sgRNAs (5’ TGTATGTGAACTCAATTAAG and 5’ AAGAAGAAGCCAATGATGTG). GFP expressing clones were sorted with a fluorescent activated cell sorter (FACS, BD FACSAria II) 48 h after nucleofection and cells were plated at a single-cell density. Clones were picked after 9 days and knockout alleles were verified by genotyping PCR.

H2B-GFP line was generated by nucleofecting the H2B-GFP plasmid (Addgene #11680) into the A17 E14Tg2a line. Randomly inserted H2B-GFP expressing cells were selected clonally and verified by live nuclear GFP expression under Nikon Perfect Focus Eclipse Ti live-cell fluorescence microscope and with FACS Aria.

The *Oct4::tdTomato Sox2::Gfp* line was generated by inserting an *IRES-dtTomato* cassette (Swanzey and Stadtfeld, 2016) into the 3’UTR of the endogenous *Oct4* locus to KH2-OKSM ESCs (Stadtfeld *et al.*, 2010) similarly as previously described (Lengner *et al.*, 2007). This was followed by heterozygous insertion of an EGFP cassette in the position of the start codon of the endogenous *Sox2* locus, using a previously described targeting vector (Ellis *et al.*, 2004).

### Cell Culture

ESCs were cultured on 0.1% gelatin (Milipore) coated dishes at 37 °C, 8% CO_2_ in ESC medium (Advanced DMEM/F12: Neurobasal (1:1) Medium (Gibco), supplemented with 2.5% mESC-grade fetal bovine serum (Corning), 1x N2 (Gibco), 1x B27 (Gibco), 2 mM L-glutamine (Gibco), 0.1 mM ß-mercaptoethanol (Gibco)) supplemented with LIF (leukemia inhibitory factor, 1,000 Uml^-1^ (Millipore)) and 2-inhibitor cocktail (3 mM CHIR (BioVision) and 1 mM PD0325901 (Sigma)) unless stated otherwise.

### Library Cloning

Synthesized oligonucleotides (Twist Bioscience) were dissolved in Buffer EB (Qiagen). 16 ng/uL single-stranded pooled oligos were amplified with NEBNext High-Fidelity 2X PCR Master Mix (NEB) with the following PCR protocol: Denaturation 98 °C for 30s, 8 cycles of 98 °C for 10s, 63 °C for 10s, 72 °C for 15s, final elongation 72 °C for 3 min. 40 μg pTF vector containing sgRNA-E+F scaffold with U6 promoter and Cas9-P2A-Zeocin with EFS-NS promoter is digested with Esp3l (Thermo Scientific) in FastDigest Buffer (10X, Thermo Scientific) and 1mM DTT in 100 uL (Chen *et al.*, 2013). The digested vector is also dephosphorylated in FastAP Thermosensitive Alkaline Phosphatase (1U/μL, Thermo Scientific) in FastDigest Buffer (10X) in 200 μL volume. Purified pooled oligos were cloned into digested pTF vector using 2X Gibson Assembly Master Mix with 10 times molar ratio of pooled oligos to digested vector and transformed to competent E.coli and plated on LB+Amp plates (Figure S1D). 565 E.coli colonies per guide ratio (>500x coverage) was used to process the 4 Maxi-Prep (Qiagen) reactions. After bacterial transformation, the plasmid library was sequenced to verify uniform sgRNA coverage and minimal bias (Figure S1E).

### Lentiviral packaging

12.5 million 293T (ATCC CRL-3216) cells were plated on 4 T225 flasks and grown in DMEM with high glucose, pyruvate (Thermo Scientific) with 10% FBS (Corning), and 2 mM L-glutamine (Gibco). When the cells were 90% confluent, 12.5 mL Opti-MEM with 630 μL Lipofectamine 3000 (Thermo Scientific) was mixed with 12.5 mL of Opti-MEM containing 61.2 μg of pMD2.G, 93.6 μg of psPAX2 and 122.4 μg of CRISPR TF library and 540 μl of P3000 enhancer (Thermo Scientific). After the mixed solution was incubated for 15 min, 6.5 mL of transfection mix is transferred per T225. 5 hours later, media containing the transfection mix was replaced with fresh DMEM with 10% FBS. 48h later, the supernatant containing lentivirus was collected from 4 T225 flasks and spun down at 1000 × g for 5 min at 4 °C to remove debris. The pooled virus-containing supernatant was filtered through a 0.45 μM PES membrane (VWR). The filtered supernatant was then spun in an ultracentrifuge for 2 hours at 20,000 g at 4 °C. 30 mL of the supernatant obtained per T225 was concentrated in 300 μL PBS1X containing 10% BSA (Sigma). Aliquots were frozen at −80 °C. To prepare individual sgRNA viruses, the above-mentioned protocol for T225 was scaled down to p100 and PEI was used to replace Lipofectamine 3000. 1:2.54 DNA to PEI ratio was used for p100.

### Mouse TF CRISPR Screen

3 × 10^6^ ESC cells were distributed to 5 12-well dishes (Thermo Scientific) with 5 × 10^5^ mESC cells per well. 4 h after cells were plated, the media was replaced with 8 μL of 100x concentrated library lentivirus mixed in 1 mL ESC medium with 1x polybrene (EMD Millipore). 16 h later, the virus solution was removed, cells were washed 1 time with PBS1X, and cells were split 1:2 to 10 12-well plates. 50 ng/μL Zeocin (Invivogen) was added to each well 24h later and transduced cells were maintained with Zeocin selection. To maintain 50-70% confluency throughout the screen, cells were transferred to 6-well plates (Thermo Fisher) on day 2 and selected for 3 additional days for a total of 5 days of selection. The first time point was obtained on day 5 post-selection. The second and third timepoints were taken on day 8 and day 12 while growing in ESC medium containing 50 ng/μL Zeocin.

We performed library readout as described previously (Chen *et al.*, 2015). Briefly, in a 15 ml conical tube, we added 6 ml of NK Lysis Buffer (50 mM Tris (Boston BioProducts), 50 mM EDTA (Ambion), 1% SDS (Invitrogen), pH 8), and 30 μl of 20 mg/ml Proteinase K (Qiagen) to 9 million cells (>500 coverage) and incubated at 55°C overnight. 30 μl of 10 mg/ml RNAse A (Qiagen, diluted in NK Lysis Buffer to 10 mg/ml and then stored at 4 °C) was added to the lysed sample, which was then inverted 25 times and incubated at 37°C for 30 minutes. Samples were cooled on ice before the addition of 2 ml of pre-chilled 7.5M ammonium acetate (Sigma) to precipitate proteins. After adding ammonium acetate, the samples were vortexed at high speed for 20 seconds and then centrifuged at ≥ 4,000 × *g* for 10 minutes. After the spin, a tight pellet was visible in each tube and the supernatant was carefully decanted into a new 15 ml conical tube. 6 ml 100% isopropanol was added to the tube, inverted 50 times, and centrifuged at ≥4,000 × *g* for 10 minutes. Genomic DNA was visible as a small white pellet in each tube. The supernatant was discarded, 6 ml of freshly prepared ice-cold 70% ethanol was added, the tube was inverted 10 times, and then centrifuged at ≥4,000 × *g* for 1 minute. The supernatant was discarded by pouring; the tube was briefly spun, and the remaining ethanol was removed using a P200 pipette. After air-drying for 10-30 minutes, the DNA changed appearance from a milky white pellet to slightly translucent. At this stage, 500 μl of 1x TE buffer (Sigma) was added, the tube was incubated at 65°C for 1 hour and then overnight at room temperature to fully resuspend the DNA. The next day, the gDNA samples were vortexed briefly. The gDNA concentration was measured using a Nanodrop (Thermo Scientific).

5 PCR1 reactions were used for 52 μg of DNA for >500x coverage. Per PCR tube, 10.4 μg of genomic DNA was used in 100 μL volume with 0.4 μL Taq-B Polymerase (Enzymatics), 10X Taq buffer (Enzymatics), 10 mM dNTPs (NEB), 10 μM Fwd (5’ GAGGGCCTATTTCCCATGATTC), and Rvs (5’ GTTGCGAAAAAGAACGTTCACGG) using the following PCR protocol: Denaturation 94 °C for 30s, 25 cycles of 94 °C for 10s, 55 °C for 30s, 68 °C for 45s, final elongation 68 °C for 2 min. For the addition of Illumina barcodes, 5 staggered forward primers and 1 reverse primer with specific barcodes containing Illumina adaptors were mixed to obtain 10 μM primer mix. 2 PCR2 reactions were used per timepoint with 5 μL of PCR1 product amplified with 25 μL Q5 High Fidelity 2X Master Mix (NEB), 5 μL of 10 μM primer mix in 50 μL volume using the following PCR protocol: Denaturation 98 °C for 30s, 10 cycles of 98 °C for 10s, 65 °C for 30s, 72 °C for 45s, final elongation 72 °C for 5 min. All samples were pooled in equimolar ratio and PCR purified pooled sample with QIAquick PCR Purification Kit (Qiagen). The purified sample was then run on a 2% E-Gel and the correct size band was extracted for PCR2 (250-270 bp) and purified with QIAquick Gel Extraction Kit (Qiagen). The pooled library was then quantified with a Low-Range Quantitative ladder on 2% E-Gel. Libraries were then sequenced on Illumina MiSeq v3 for 150 cycles single end at the genomics core facility at NYU.

Sequences flanking the 20 nucleotide sgRNA sequence were trimmed using cutadapt. The remaining 20 nucleotide sgRNA sequence reads were aligned to a library index generated by 17820 sgRNA sequences using bowtie. Each sgRNA ratio was calculated by normalizing to the total reads captured per timepoint to be used for downstream analysis.

### Human TF CRISPR screen

NYGCe001-A human embryonic stem cells (derived from HUES66) were used for the human TF CRISPR screen (Lu and Sanjana, 2019). NYGCe001-A cells were maintained using the Enhanced Culture Platform (ECP), as described previously by Cowan and colleagues (Schinzel *et al.*, 2011). Briefly, hESCs were cultured in Essential 8 media (Thermo Fisher) supplemented with 100 μg/mL Normocin (InvivoGen) and cultured in standard tissue culture dishes coated with Geltrex (Thermo Fisher) at 37 °C in 5% CO2. Accutase (STEMCELL) was used for passaging the cells. 10 μM Rho kinase inhibitor (ROCKi) Y-27632 (MilliporeSigma) was added to the culture medium at each passage. ROCKi was removed at the subsequent media change (typically 24 hours later).

For the Human TF CRISPR-Cas9 library, we designed sgRNAs to target 1891 known human TFs using the GUIDES web tool (http://guides.sanjanalab.org) with 10 sgRNAs per TF and also included 1000 non-targeting (negative control) sgRNAs (Ravasi *et al.*, 2010; Meier, Zhang and Sanjana, 2017). The library was cloned and packaged using the same methods described above (Library cloning and Lentiviral packaging), except that no BSA was added to concentrated lentivirus.

Briefly, 80 × 10^6^ NYGCe001-A cells were transduced with human TF CRISPR-Cas9 library lentivirus. 2 days after transduction, 4 μg/μL Zeocin (Invivogen) were added to culture media, and then cells were maintained in selection media for 2 days. After selection, cells were cultured for 24 days, then harvested for DNA extraction and library amplification (see DNA extraction and library amplification above). Cells were passaged whenever they reached approximately 75% confluence. After each passage, at least 10 × 10^6^ cells were maintained in culture to ensure ~500-fold coverage of the total number of sgRNAs in the Human TF CRISPR library.

To identify depleted guide RNAs, we compared sgRNA representation from Day 24 post-selection to the plasmid library. To identify significantly depleted guides, we assessed how many sgRNAs for each TF were depleted below the 5th percentile of non-targeting guide RNAs and also performed robust rank aggregation (RRA) analysis.

### Competition experiment with qPCR

U6-sgRNA-E+F scaffold and EFS-NS-Cas9-P2A-ZeocinR-WPRE digested from pTF vector with AflII and HindIII and cloned into the piggyBac backbone (De Santis *et al.*, 2018). sgRNAs targeting *Klf5* (5’ TGGCGAATTAACTGGCAGAG), *Nanog* (5’ TGTCCTTGAGTGCACACAGC), *Oct4* (5’ CCGCCCGCATACGAGTTCTG), *Zbtb10* (5’ AGAAACGGCTGCCTGCAACC), *Zbtb11* (5’ AGCGCACAAGTCTGTCCTCT), *Zfp131* (5’ GTTCTTTAAAGTGTCCAAAG) and non-targeting sgRNA (5’ GCCGCAACGTTAGATGTATA) were cloned into piggyBac plasmids. Equimolar ratio (0.5 μg) of plasmids were pooled and co-nucleofected (Lonza) with 0.5 μg pBac transposase to mESCs (De Santis *et al.*, 2018). DNA was collected with >10000 cells per sgRNA coverage d1, d3, d6, d9, and d13 posttransduction. PCR1 reaction described above was used to amplify the sgRNA construct from the genome. qPCR primers were designed to overlap sgRNA and scaffold (5’ CGGTGCCACTTTTTCAAGTTG). 10 μL Maxima SYBR Green brilliant PCR amplification kit (Thermo Scientific), 5 uL forward and reverse primer mix (2 nM), 5 μL of PCR1 reaction (2 ng) were combined for the qPCR reaction using CFX 96 Touch Biorad qPCR thermocycler (Biorad). Δct was calculated by subtracting the mean Cq of each sgRNA to a non-targeting control. ΔΔct was calculated by subtracting the Δct of each timepoint from the d1 original abundance. The average depletion rate was drawn with error bars (n=3).

### *Oct4::TdTomato Sox2::Gfp* experiments

5×10^5^ *Oct4::tdTomato Sox2::Gfp* cells were plated at a single-cell density to 6-well plates and after attachment to the plate, transduced with 36 sgRNAs (3 positive controls (*Klf5, Nanog, Oct4*), 2 nontargeting negative controls and 7 candidate TFs). Transduced cells were selected on 50 ng/μL zeocin for 5 days post-transduction. ESCs were cultured in +2i+LIF media conditions for the initial 4 days and media conditions switched to −2i-LIF for 24h before proceeded to FACS.

### Immunocytochemistry

Embryonic stem cells were fixed with 4% paraformaldehyde in 1X PBS for 10 mins, washed 3 times with 1X PBS, treated with 0.02% Triton-X (Thermo Fisher) for 10 mins and then with blocking buffer for 30 mins (10% FBS, 0.05% NaAz in 1X PBS), stained with primary antibody overnight at 4°C, washed 3 times with 1X PBS, stained with secondary antibody at room temperature for 1h, washed 3 times with 1X PBS and mounted with Fluoroshield with DAPI (Sigma). Images were acquired with a Nikon Perfect Focus Eclipse Ti live-cell fluorescence microscope. We used antibodies to V5 (R960-25, Thermo Fisher Scientific; 1:5000), OCT3/4 (Sc-5279, Santa Cruz; 2mg/ml), GFP (ab13970, Abcam; 1:5000), HA (ab9110, Abcam; 1:5000), Alexa 488 (A-11029, A-11015), Alexa 568 (A-11036) secondary antibodies were used (Thermo Fisher, 1:2000).

### Competition experiments with H2B-GFP

10^6^ H2B-GFP cells were co-plated with 10^6^ E14Tg2a wt, i*Zbtb10*-HA *Zbtb10Δ, iZbtb11-HA Zbtb11Δ* and *iZfp131-HA Zfp131Δ* cells on 6-well plates and grown in +2i+LIF conditions without doxycycline. Cells were fixed d1 and d5 after plated and stained for GFP to calculate GFP/DAPI ratio. Due to plating error, d1 values were normalized to 1:1 and d5 values were calculated according to the normalization. Competition rates were calculated with error bars (n=3).

### RNA-Seq

Cells were collected after grown for 3d in +2i or −2i conditions in the absence of doxycycline. TRIzol LS Reagent (LifeTechnologies) was used to extract RNA and RNAeasy mini kit (Qiagen) and used to purify RNA. Agilent High Sensitivity RNA Screentape (Agilent) was used to determine RNA integrity. 500 ng RNA from cells that was spiked-in with ERCC Exfold Spike-in mixes (Thermo Fisher, 4456739) was used for the generation of RNA-seq libraries. TruSeq Stranded mRNA Library Preparation kit (Illumina, 20020594) was used to prepare RNA-seq libraries. High Sensitivity DNA ScreenTape (Agilent, 5067-5584) was used to verify library sizes. A KAPA library amplification kit was used on Roche Lightcycler 480 to quantify the library prior to sequencing. Libraries were sequenced on Illumina NextSeq 500 using V2.5 chemistry (75 cycles, single-end 75bp) at the Genomics Core Facility at NYU.

### ChIPseq

Wildtype and mutant cells were collected after being grown for 3d in +2i or −2i conditions in the presence or absence of 3 ng/uL doxycycline. 1 mM DSG (ProteoChem) was used for crosslinking at RT for 15min. 1% Formaldehyde (vol/vol) was added for an additional 15min until quenched with glycine (Sigma). Cells were separated into aliquots of 25×10^6^ and centrifuged to freeze pellets at −80°C. Thawed cells were lysed in 5mL of 50 mM HEPES-KOH pH 7.5 (Gibco), 140 mM NaCl (Thermo Fisher), 1 mM EDTA pH 8.0, 10% glycerol (vol/vol) (Thermo Fisher), 0.5% Igepal (vol/vol) (Sigma), 0.25% Triton X-100 (vol/vol) with 1×protease inhibitors (Roche) for 10min at 4°C. Cells were centrifuged for 5min at 1100 rpm and resuspended in 5mL of 10 mM Tris-HCl pH 8.0 (Boston BioProducts), 200 mM NaCl, 1 mM EDTA pH 8.0, 0.5 mM EGTA pH 8.0 (Boston BioProducts) with 1×protease inhibitors, and incubated for an additional 10 min at 4°C on a rotating platform. Cells were centrifuged for 5min at 1100 rpm and resuspended in 2mL of Sonication Buffer (50 mM Hepes pH 7.5, 140 mM NaCl, 1 mM EDTA pH 8.0, 1 mM EGTA pH 8.0, 1% Triton X-100 (vol/vol), 0.1% sodium deoxycholate (wt/vol), 0.1% SDS (vol/vol) with 1×protease inhibitors). Sonication was performed in two Bioruptor tubes per sample with 0.45 g sonication beads. Bioruptor Pico (Diagenode) was used for 18 cycles of 30 sec on and 30 sec off to sonicate to an average size of approximately 200 bp. To immunoprecipitate, Dynabeads protein-G (Thermo Fisher) and antibodies (HA, H3K4me3, H3K27me3, H3K27ac, SIN3A, CHD4) were incubated with sonicated chromatin for 16h at 4°C on a rotating platform. After the immunoprecipitation, the sample was washed with sonication buffer, high salt sonication buffer (500 mM NaCl, LiCl wash buffer (20 mM Tris-HCl pH 8.0, 1 mM EDTA pH 8.0, 250 mM LiCl (Sigma), 0.5% Igepal (vol/vol), 0.5% sodium deoxycholate (wt/vol)), and TE buffer (10 mM Tris-HCl pH 8.0, 1 mM EDTA pH 8). The sample was eluted in elution buffer (50 mM Tris-HCl pH 8.0, 10 mM EDTA pH 8.0, 1% SDS (vol/vol)) by incubating for 45min at 65°C. Crosslinks were reversed by incubating the sample for 16h at 65°C. RNA was removed by 0.2 mg/mL RNAse A (Sigma) in 200 μL of TE for 2h at 37°C. To remove protein, 0.2 mg/mL Proteinase K (Invitrogen) and CaCl_2_ (Sigma) were added and the sample incubated at 55°C for 30min. DNA was first purified with phenol:chloroform:isoamyl alcohol (25:24:1; vol/vol) (Invitrogen) and then by performing salt-ethanol precipitation. DNA pellets were resuspended in 50 uL H_2_O. Illumina DNA sequencing libraries were prepared with half of the ChIP sample and a 1:100 dilution of the input sample in H_2_O. Library preparation was performed with end repair, A-tailing, and ligating multiplexed adapters (Illumina-compatible Bioo Scientific). Agencourt AmpureXP beads (Beckman Coulter) were used to remove unligated adapters. Libraries were then amplified by PCR with TruSeq primers (Sigma) and Phusion polymerase (New England Biolabs). Libraries with sizes ranging between 250-550bp were gel purified (Qiagen). KAPA library amplification kit was used on Roche Lightcycler 480 to quantify the library prior to sequencing. Libraries were sequenced on Illumina NextSeq 500 using V2.5 chemistry (75 cycles, single-end 75bp) at the Genomics Core Facility at NYU.

### scRNAseq

*Zbtb11Δ* and *Zfp131Δ* cells were mixed with H2B-GFP expressing wt cells in a 9:1 ratio to have 1000 cells/ul. CellTrics 30 μM (Cat #04–004-2326) was used to remove debris and clumps. 10X Genomics Chromium Single Cell 3’ library kit was used to generate the single-cell library with a cell recovery rate of 10.000 cells (120262 Chromium™ i7 Multiplex Kit, 120236 Chromium™ Single Cell 3’ Chip Kit v2, 120237 Chromium™ Single Cell 3’ Library & Gel Bead Kit v2). Agilent High Sensitivity DNA D1000 Screentape (5067–5585) was used to detect fragment length distribution of the library. KAPA library amplification kit was used on Roche Lightcycler 480 to quantify the library prior to sequencing. Libraries were sequenced on Illumina NovaSeq 6000 using SP chemistry (100 cycles, 26×98 bp) at the Genomics Core Facility at NYU.

### Candidate TF identification with False Discovery Rate

In the False Discovery Rate method, for each time point, normalized counts of each sgRNA were calculated. Log2 scores were then calculated and ranked by taking the log2 of the final count divided by the initial count. We set the thresholds for calling depleted and enriched targeting sgRNAs setting an empirical <0.05 false discovery rate (FDR). Any targeting sgRNA that was underrepresented less than 50^th^ most underrepresented non-targeting sgRNA was determined to be a depleted sgRNA with FDR <0.05 (marked by orange in Figure 1C, S2C). Conversely, any sgRNA that was overrepresented more than the 50^th^ most overrepresented non-targeting sgRNA was determined to be an enriched sgRNA with FDR<0.05 (marked by blue in Figure 1C, S2C). Depleting and enriching sgRNAs were then appointed to genes to determine how many sgRNAs were depleted or enriched per each gene.

### Candidate TF identification with Linear Model

After normalization, a mixed linear model (Gelman and Hill 2006) was used to evaluate guide enrichment or depletion. The model has the following formula:

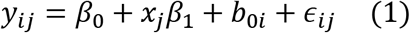

Where *y_ij_* is the log counts of guide *i*. at the *j*-th time point; *β*_0_ is the overall intercept; *β*_1_ is the overall slope for time; *x_j_* is the *j*-th time point; *b_0i_* is the random intercept for guide *i*. and *ϵ_j_* is the random error. A random intercept model was chosen to account for the variability in the guides’ initial observed counts. Hypothesis testing was done using the likelihood ratio test via ANOVA (Kaufmann and Schering, 2014). The model was compared to a reduced version that does not account for time. P-value adjustment was performed using the Holm (Sture Holm 1979) method. An adjusted p-value < 0.05 was used to identify candidates.

To take advantage of the time course data captured throughout our experiment, the linear model was modified to classify each of the hits as early, late, or a stably depleting gene. In this new model, the time parameter was encoded using the following factors:

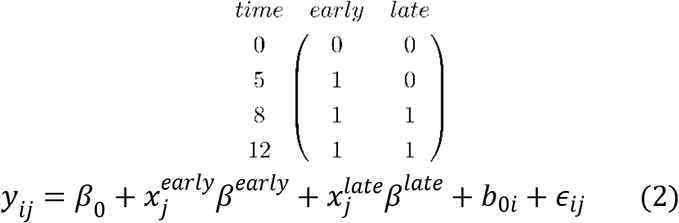

Hypothesis testing was performed as described above. Depending on the significance of each time factor, our genes were classified as follows:

| $time^{early}$ | $time^{late}$ | Classification |
| --- | --- | --- |
| True | True/False | Early |
| False | False | Stable |
| False | True | Late |

“Early depleting” genes include core cellular machinery and the core PGRN TFs *Oct4* and *Nanog* (Figure S4B-D). “Stable and late depleting” genes include cellular stress and DNA repair factors along with accessory TFs in the PGRN, such as *Tfcp2l1* (“stable depleting”), *Stat3* (“late depleting”), and *Nr0b1* (“late depleting”) (Figure S4B, E-G). Accessory TFs do not rapidly deplete due to the time required for their mutations to exert an effect on the levels of core TFs. The screen depth and time series were sensitive enough to identify sgRNAs targeting partially redundant genes like *Klf2* and *Klf5* as “late depleting” TFs (Figure S4H, I).

### RNAseq Data Processing

RNA-seq fastq files were aligned to the mouse genome (version mm10) using Hisat2 (version 2.1.0) (Kim, Langmead and Salzberg, 2015). FeatureCounts R package was used to assign reads to genomic features (Liao, Smyth and Shi, 2014). The DESeq2 package was used for differential gene expression analysis and data visualization (volcano plots, heatmaps, PCA plots)(Love, Huber and Anders, 2014). A q-value cutoff of < 0.01 was used for significantly misregulated genes. PANTHER (version 14) (http://geneontology.org) was used to perform Gene Ontology term enrichment analysis (Mi *et al.*, 2019).

### scRNAseq Data Processing

Fastq files were generated by using 10X Genomics CellRanger (version 4.0.0) with default settings. New fasta and annotation files were created using mkref in mm10 for the detection of the H2B::GFP sequence (Addgene #11680). CellRanger count function was used to assign reads to genomic features. 7510 cells were estimated for *Zbtb11Δ* and H2B-GFP population whereas 6982 cells were estimated for *Zfp131Δ* and H2B-GFP population. Seurat R package (version 3.0) was used for differential gene expression analysis and data visualization (UMI plots, violin plots, gene-specific UMI plots)(Stuart *et al.*, 2019).

### ChIP-seq Data Processing and Differential Analysis

Fastq files were aligned to the mouse genome (mm10) using Bowtie (v1.0.1) with options “-q-best-strata -m 1-chunkmbs 1024”. MultiGPS (v.0.74) was used to call peaks after alignment on all transcription factors. A q-value cutoff of <0.01 was used to identify significant binding events. Histone modification and ATAC-seq peaks were called using the DomainFinder module in SeqCode (https://github.com/seqcode/seqcode-core/blob/master/src/org/seqcode/projects/seed/DomainFinder.java). Contiguous 50-bp genomic bins with significantly higher read enrichment compared with normalized input were identified (binomial test, p <0.05). Differential binding and modification analyses were done using DESeq2 (v1.28.1) with an adjusted p-value cutoff of <0.05 and all default options. To create the count matrix for DESeq2, peaks were first called independently in the wildtype and knockout ChIP-seq datasets. Overlapping peak regions were merged between the wt and KO. Peak regions found exclusively in either file were also included. The final count matrix was created using Bedtools coverage with the “-counts” option separating each replicate into its count column. Before running DESeq2 on the matrix, counts were normalized by the length of the corresponding region for histone modifications (domains have differing lengths).

### Motif Finding

For motif finding, the top 1000 peaks were selected from each zinc finger ChIP-seq dataset based on MultiGPS q-value. Following this, bedtools getFasta was used to retrieve the sequences underlying these binding events. Sequences were fed through RepeatMasker (v4.0.7) on default parameters. Finally, meme-chip from the MEME suite (v4.11.3) was performed on the masked sequences using the options “-meme-nmotifs 5 -meme-mod zoops -meme-minw 6 -meme-maxw 20”.

### Data Visualization for Heatmaps and Profile Plots

Heatmaps and profile plots were generated using Deeptools (v3.1.3). Bigwigs were created directly from the BAM files using RPKM normalization within Deeptools bamCoverage (--binSize 50 -- normalizeUsing RPKM). Other normalizations were also tested. Using RPKM normalization kept the noise the same visually between replicates and conditions when plotting 5000 random regions in the genome (for within-study datasets). Replicates were concatenated together before creating the bigwig file as suggested by the Deeptools manual. Matrices for the heatmaps and profile plots were created using Deeptools computeMatrix with the options “reference-point --referencePoint center -a 1000 -b 1000”. Heatmaps themselves were generated using Deeptools plotHeatmap with all default options outside of changing sample colors. Max values were chosen by comparing 5000 random noise regions in each sample for within-study data and inspecting these visually. The datasets retrieved from other studies had significantly more noise than from our own. Profile plots were created using Deeptools plotProfile with the option “--perGroup”.

### Overlap Analysis for ChIPseq Data

Overlap percentages were calculated using bedtools intersect with the “-u” option. The first file or “-a” was always the zinc finger. In other words, the percentages represent the number of zinc finger peaks (A) that intersect the other transcription factor or histone modification regions (B) in question. For figure 5A, ZBTB11 was used as “-a.” Observed/expected values in Figure 5B were calculated using a python script. In brief, this script created bed files that had the same number of regions and lengths of the regions within the second bed file (B) and intersected these with the zinc finger bed files (A). The script discounted blacklist regions to increase accuracy. This was performed 30 times to get the average number of times one would expect file A to overlap with file B. The mean was used as expected in Figure 5 and the actual value from our data was used as the observed.

