## Supplementary Figures for "The BTB transcription factors ZBTB11 and ZFP131 maintain pluripotency by pausing POL II at pro-differentiation genes"

Figure S1

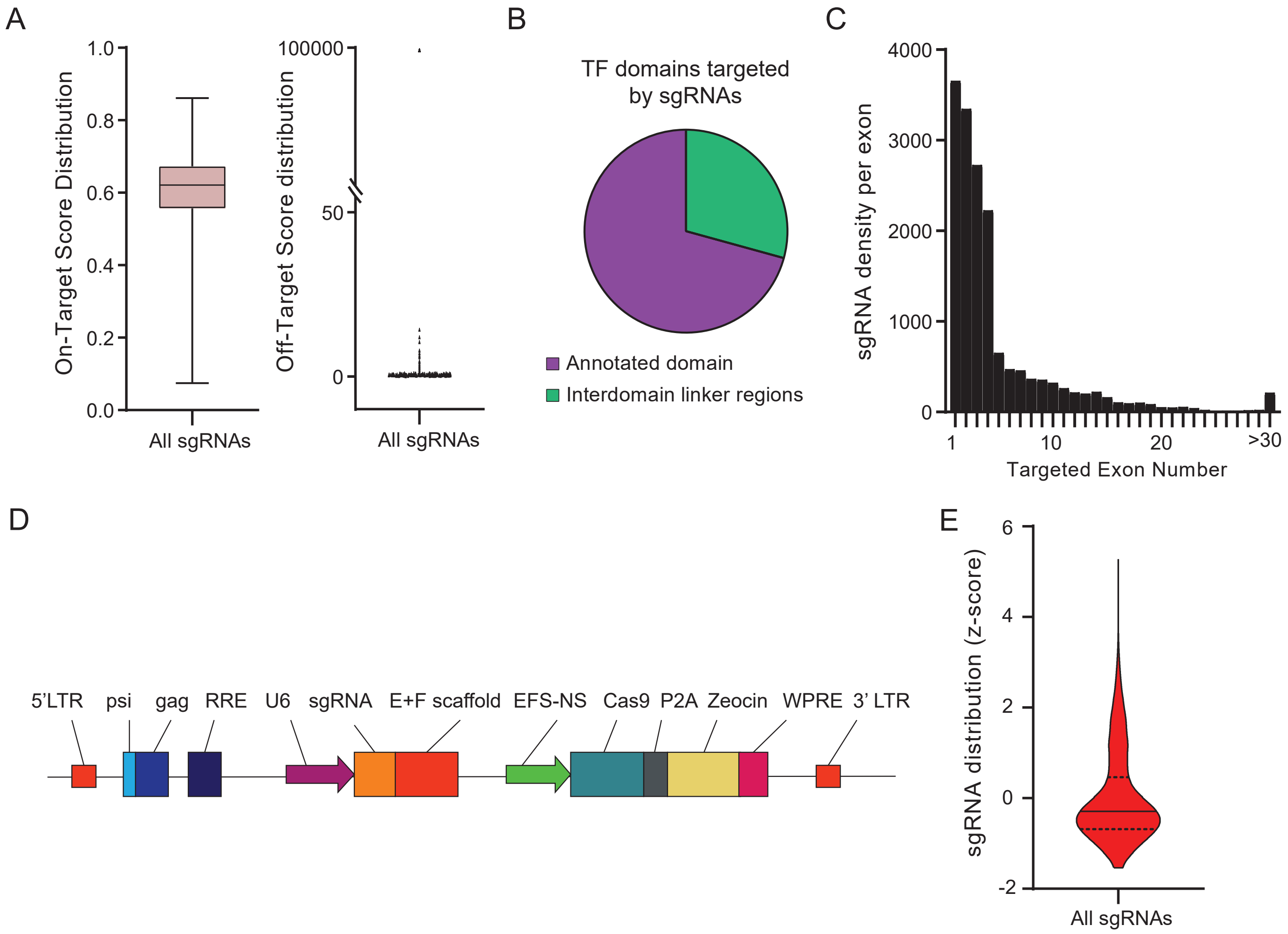

A
1.0
0.8
0.6
0.4
0.2
0.0
On-Target Score Distribution
All sgRNAs
100000
50
0
Off-Target Score distribution
All sgRNAs
B
TF domains targeted by sgRNAs
Annotated domain
Interdomain linker regions
C
4000
3000
2000
1000
0
sgRNA density per exon
1
10
20
>30
Targeted Exon Number
D
5'LTR
psi
gag
RRE
U6
sgRNA
E+F scaffold
EFS-NS
Cas9
P2A
Zeocin
WPRE
3' LTR
E
6
4
2
0
-2
sgRNA distribution (z-score)
All sgRNAs

**Figure S1. CRISPR TF library targets 1682 TFs in the mouse genome.**

**(A)** On-target and off-target score distribution of all sgRNAs in the library based on the GUIDES algorithm.

**(B)** Percentage of targeted annotated TF domains in the library based on Pfam. 71% of the sgRNAs target an annotated functional domain.

**(C)** Exon targeting density of sgRNAs in the library. sgRNAs were designed to target 5' exons to increase the chance of generating a frameshift.

**(D)** The lentiviral expression vector for sgRNA and Cas9. psi+, Psi packaging signal; RRE, Rev response element; cPPT, central polypurine tract; EFS, elongation factor 1a short promoter; Flag, Flag octapeptide tag; P2A, 2A self-cleaving peptide; Zeo, zeocin resistance gene; WPRE, post-transcriptional regulatory element; EF1a, elongation factor 1a promoter.

**(E)** Representation of the normalized number of reads of sgRNAs in the plasmid library with Z-score distribution. The solid line represents the mean z-score. Dashed lines represent quartiles.

#### Figure S2

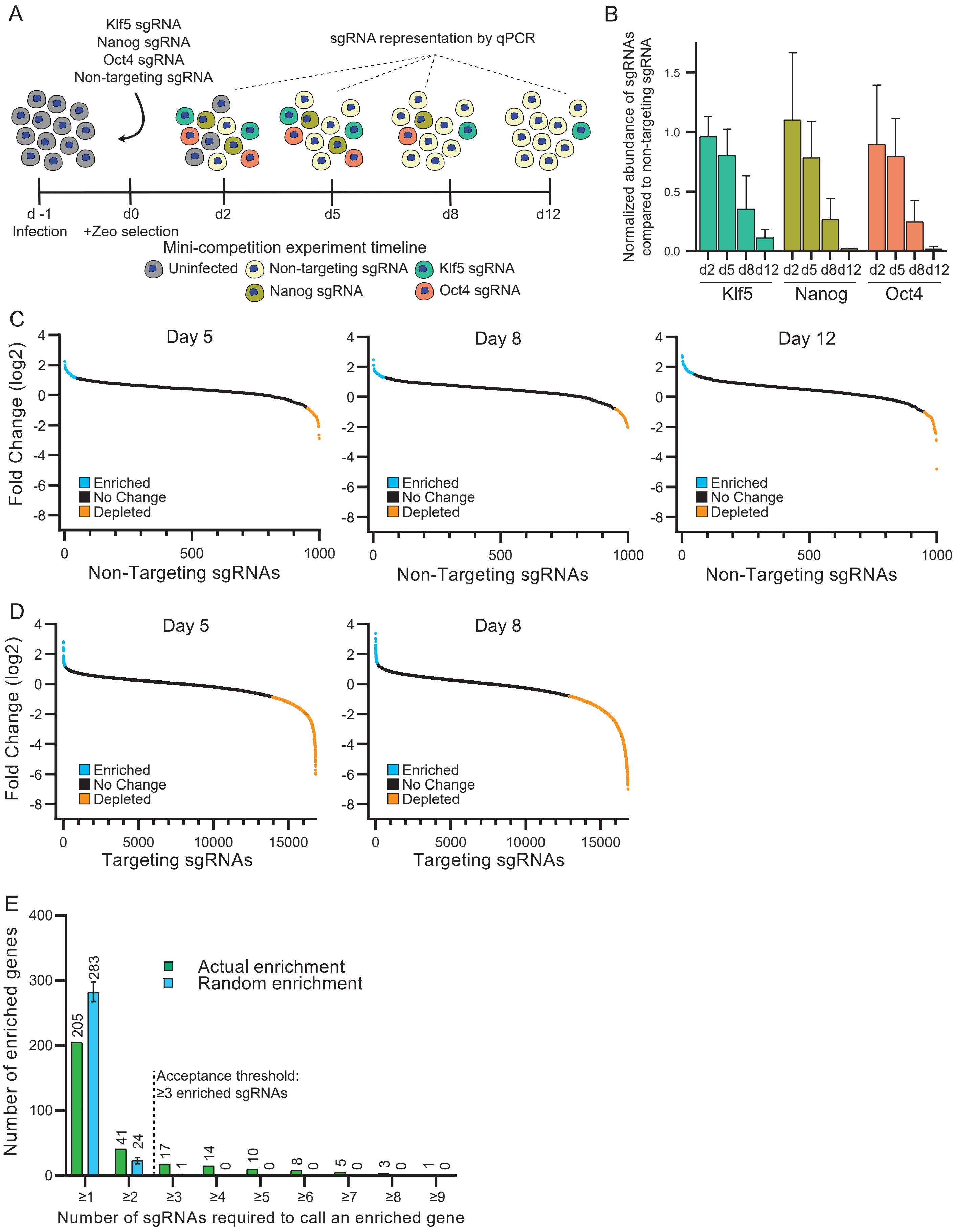

A
Klf5 sgRNA
Nanog sgRNA
Oct4 sgRNA
Non-targeting sgRNA
sgRNA representation by qPCR
d -1
Infection
d0
+Zeo selection
d2
d5
d8
d12
Mini-competition experiment timeline
Uninfected
Non-targeting sgRNA
Klf5 sgRNA
Nanog sgRNA
Oct4 sgRNA
B
Normalized abundance of sgRNAs compared to non-targeting sgRNA
1.5
1.0
0.5
0.0
d2 d5 d8 d12 d2 d5 d8 d12 d2 d5 d8 d12
Klf5
Nanog
Oct4
C
Day 5
Day 8
Day 12
Fold Change (log2)
Enriched
No Change
Depleted
Non-Targeting sgRNAs
D
Day 5
Day 8
Fold Change (log2)
Enriched
No Change
Depleted
Targeting sgRNAs
E
Number of enriched genes
Actual enrichment
Random enrichment
205
283
41
24
17
1
14
0
10
0
8
0
5
0
3
0
1
0
Acceptance threshold:
≥3 enriched sgRNAs
≥1
≥2
≥3
≥4
≥5
≥6
≥7
≥8
≥9
Number of sgRNAs required to call an enriched gene

**Figure S2. Determination of candidate factors based on the false discovery rate from non-targeting sgRNAs.**

**(A)** Schematic overview of the qPCR competition assay to identify depletion dynamics. Representation of non-targeting and *Klf5*, *Oct4*, and *Nanog* targeting sgRNAs infected cells over 12 days of culture.

**(B)** Normalized abundance of *Klf5*, *Oct4*, *Nanog* targeting sgRNAs compared to non-targeting sgRNA with qPCR in collected samples for each time point (n=2). The abundance of pluripotency TFs is nullified within 12d.

**(C)** Log2 fold change of non-targeting sgRNAs on day 5, day 8, day 12 compared to the initial library representation. 50 most enriched sgRNAs are in blue and 50 most depleted sgRNAs are in orange. $50^{th}$ most enriched and $50^{th}$ most depleted sgRNAs were selected as the thresholds to call depleted and enriched targeting sgRNAs for each timepoint.

**(D)** Log2 fold change of targeting sgRNAs on day 5 and day 8 compared to the initial library representation. sgRNAs depleted more than the $50^{th}$ most depleted non-targeting sgRNA are in orange and enriched more than the $50^{th}$ most enriched non-targeting sgRNA are in blue.

**(E)** The number of actual sgRNAs enriched and mean sgRNAs number with random enrichment calculated by 1000 permutation tests with a different number of sgRNA per gene thresholds. TFs with more than or equal to 3 enriched sgRNAs are unlikely to be false positives.

Figure S3

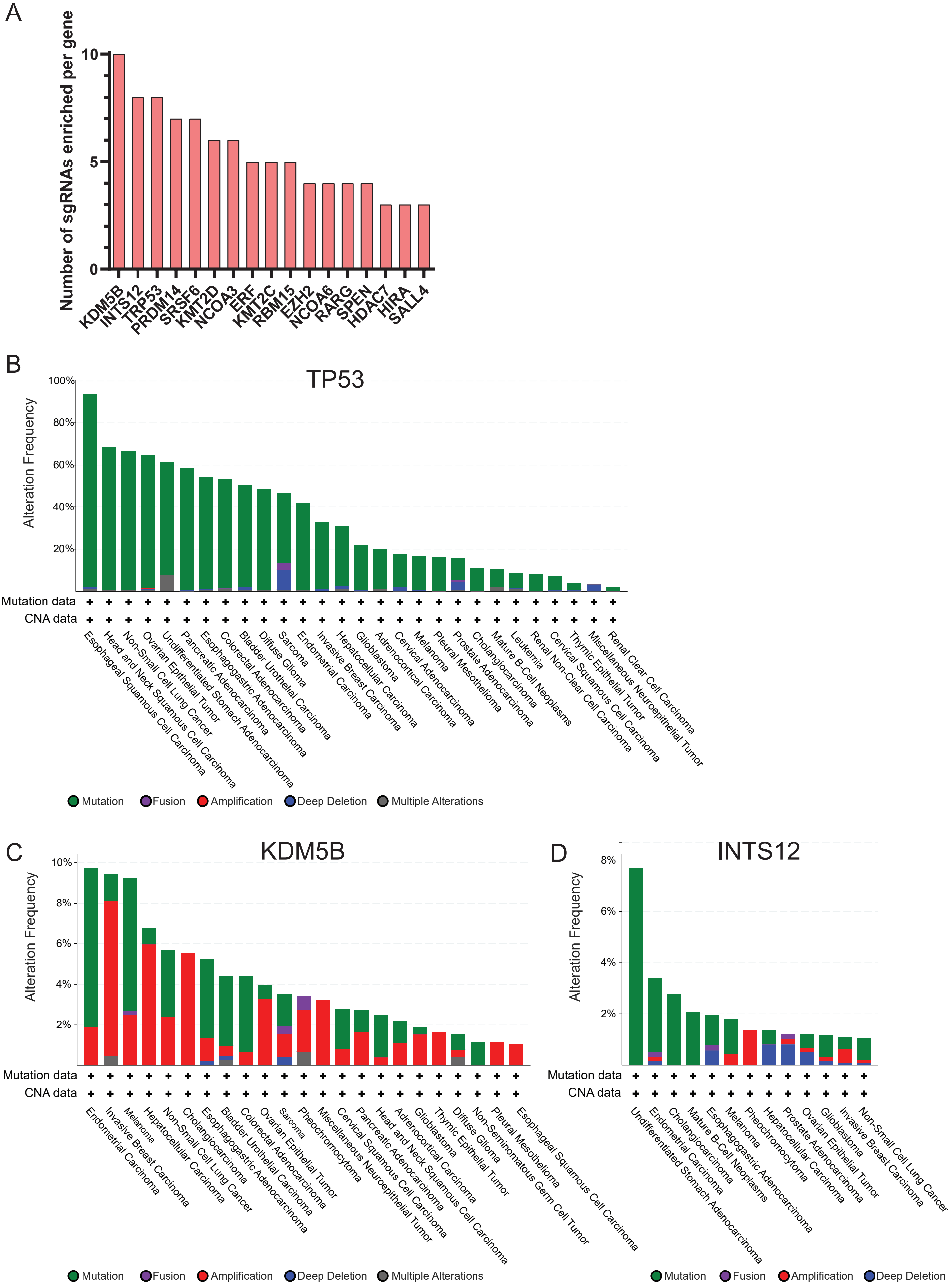
A
Number of sgRNAs enriched per gene
0
5
10
KDM5B
INTS12
TRP53
PRDM14
SRSF6
KMT2D
NCOA3
ERF
KMT2C
RBM15
EZH2
NCOA6
RARG
SPEN
HDAC7
HIRA
SALL4
B
TP53
Alteration Frequency
100%
80%
60%
40%
20%
Mutation data
CNA data
Esophageal Squamous Cell Carcinoma
Head and Neck Squamous Cell Carcinoma
Non-Small Cell Lung Cancer
Ovarian Epithelial Tumor
Undifferentiated Stomach Adenocarcinoma
Pancreatic Adenocarcinoma
Esophagogastric Adenocarcinoma
Colorectal Adenocarcinoma
Bladder Urothelial Carcinoma
Diffuse Glioma
Sarcoma
Endometrial Carcinoma
Invasive Breast Carcinoma
Hepatocellular Carcinoma
Glioblastoma
Adrenocortical Carcinoma
Cervical Adenocarcinoma
Melanoma
Pleural Mesothelioma
Prostate Adenocarcinoma
Cholangiocarcinoma
Mature B-Cell Neoplasms
Leukemia
Renal Non-Clear Cell Carcinoma
Cervical Squamous Cell Carcinoma
Thymic Epithelial Tumor
Miscellaneous Neuroepithelial Tumor
Renal Clear Cell Carcinoma
Mutation
Fusion
Amplification
Deep Deletion
Multiple Alterations
C
KDM5B
Alteration Frequency
10%
8%
6%
4%
2%
Mutation data
CNA data
Endometrial Carcinoma
Invasive Breast Carcinoma
Melanoma
Hepatocellular Carcinoma
Non-Small Cell Lung Cancer
Cholangiocarcinoma
Esophagogastric Adenocarcinoma
Bladder Urothelial Carcinoma
Colorectal Adenocarcinoma
Ovarian Epithelial Tumor
Sarcoma
Pheochromocytoma
Miscellaneous Neuroepithelial Tumor
Cervical Squamous Cell Carcinoma
Pancreatic Adenocarcinoma
Head and Neck Squamous Cell Carcinoma
Adrenocortical Carcinoma
Glioblastoma
Thymic Epithelial Tumor
Diffuse Glioma
Non-Seminomatous Germ Cell Tumor
Pleural Mesothelioma
Esophageal Squamous Cell Carcinoma
Mutation
Fusion
Amplification
Deep Deletion
Multiple Alterations
D
INTS12
Alteration Frequency
8%
6%
4%
2%
Mutation data
CNA data
Undifferentiated Stomach Adenocarcinoma
Endometrial Carcinoma
Cholangiocarcinoma
Mature B-Cell Neoplasms
Esophagogastric Adenocarcinoma
Melanoma
Pheochromocytoma
Hepatocellular Carcinoma
Prostate Adenocarcinoma
Ovarian Epithelial Tumor
Glioblastoma
Invasive Breast Carcinoma
Non-Small Cell Lung Cancer
Mutation
Fusion
Amplification
Deep Deletion

**Figure S3. Enriched genes are mutated in cancers.**

**(A)** 17 genes have 3 or more sgRNAs enriched on day 12.

**(B)** *Tp53*, **(C)** *Kdm2b*, **(D)** *Ints12* mutations incidence in various types of cancers (51799 patients). Data obtained from cBioPortal for Cancer Genomics. Enriched sgRNAs are targeting genes that are growth limiting, which are also found in cancer types.

### Figure S4

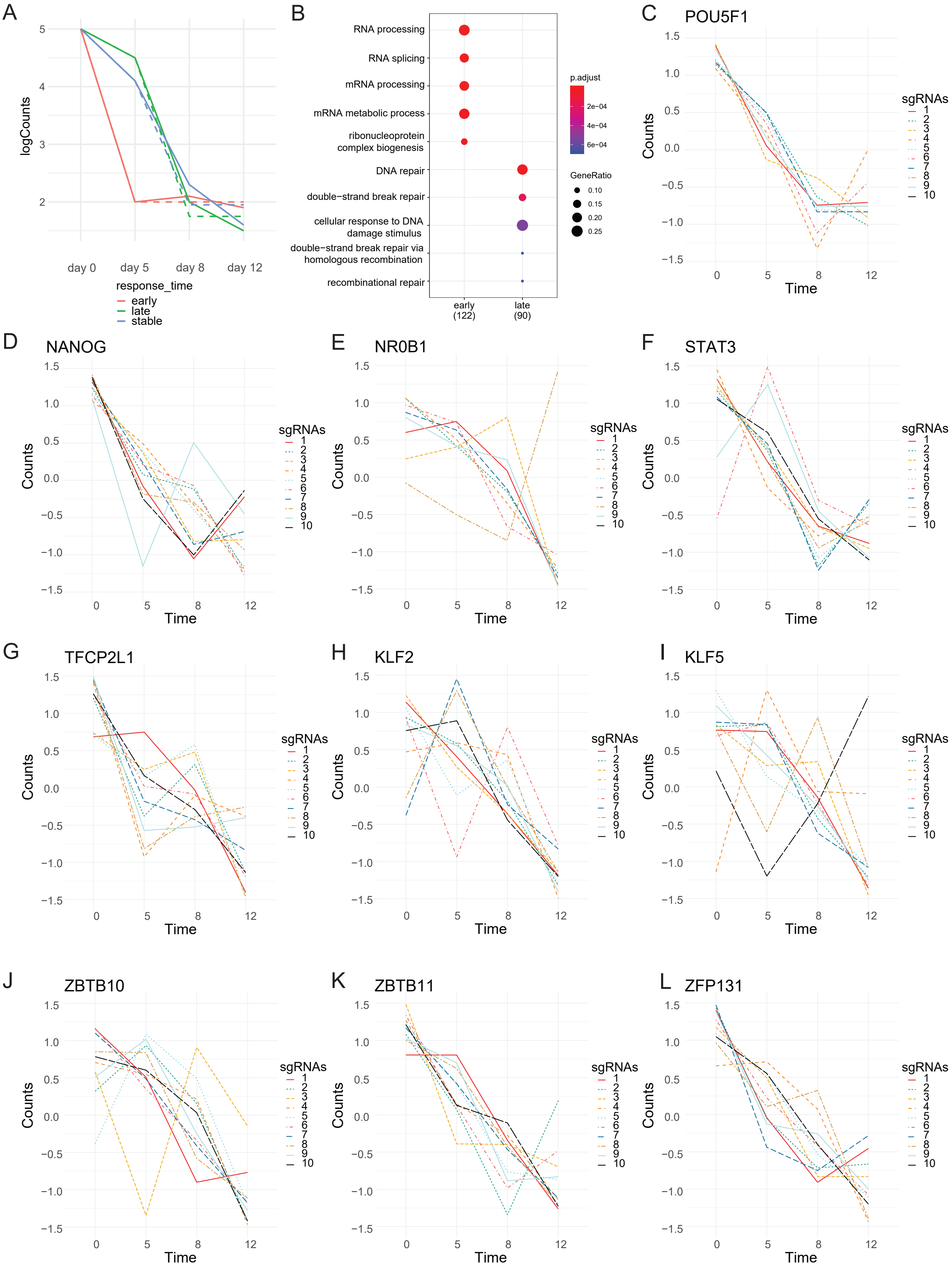

A
logCounts
day 0
day 5
day 8
day 12
response_time
early
late
stable
B
RNA processing
RNA splicing
mRNA processing
mRNA metabolic process
ribonucleoprotein complex biogenesis
DNA repair
double−strand break repair
cellular response to DNA damage stimulus
double−strand break repair via homologous recombination
recombinational repair
early (122)
late (90)
p.adjust
2e−04
4e−04
6e−04
GeneRatio
0.10
0.15
0.20
0.25
C
POU5F1
D
NANOG
E
NR0B1
F
STAT3
G
TFCP2L1
H
KLF2
I
KLF5
J
ZBTB10
K
ZBTB11
L
ZFP131
Counts
Time
sgRNAs

**Figure S4. Determination of candidate factors using a linear modeling method.**

**(A)** Linear modeling for differential depletion categories of genes based on normalized sgRNA log counts. Three categories: early, stable, and late depleting were identified. The solid line is how the linear modeling is predicted and the dashed line is how the data fit to the projection.

**(B)** GO-enrichment analysis of early (n=122) and late (n=90) depleting factors throughout the screen. Gene Ratio is calculated with the number of occurrences in specific GO categories divided by the total amount of depleted TFs per category. Early category contains genes that are in core transcription machinery whereas late category contains genes that are playing roles in DNA damage.

**(C)** Linear regression of 10 sgRNAs targeting known core pluripotency factors *Oct4*, **(D)** *Nanog*, accessory factors (**(E)** *Nr0b1*, **(F)** *Stat3*, **(G)** *Tfcp2l1*), redundant accessory factors (**(H)** *Klf2*, **(I)** *Klf5*) and candidate hits from the screen (**(J)** *Zbtb10*, **(K)** *Zbtb11*, **(L)** *Zfp131*). *Oct4*, *Nanog*, *Zbtb11,* and *Zfp131* are in the early depleting category whereas accessory factors and *Zbtb10* are in the stable and the late depleting categories.

Figure S5

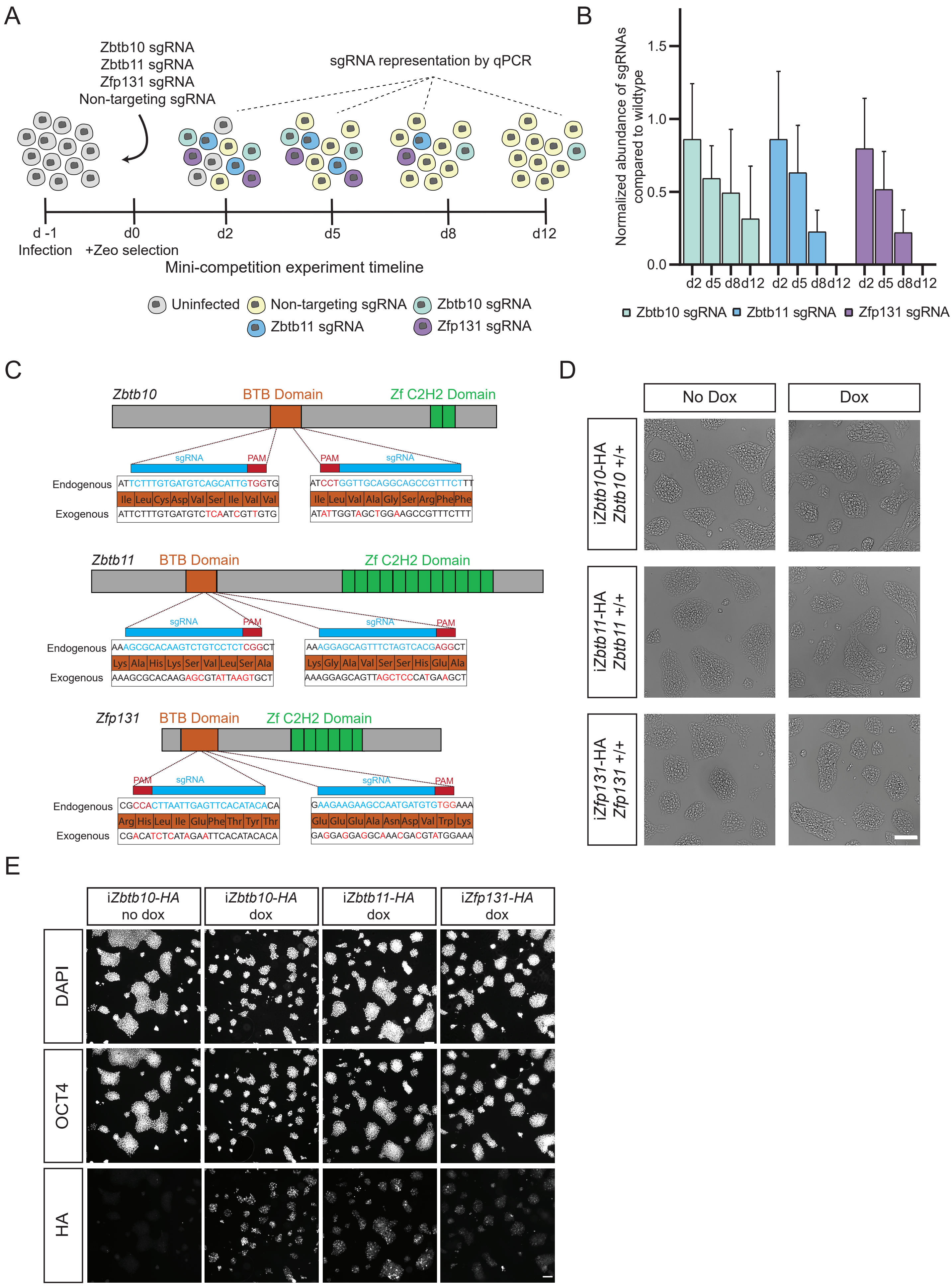
A
Zbtb10 sgRNA
Zbtb11 sgRNA
Zfp131 sgRNA
Non-targeting sgRNA
sgRNA representation by qPCR
d -1
Infection
d0
+Zeo selection
d2
d5
d8
d12
Mini-competition experiment timeline
Uninfected
Non-targeting sgRNA
Zbtb10 sgRNA
Zbtb11 sgRNA
Zfp131 sgRNA
B
Normalized abundance of sgRNAs compared to wildtype
1.5
1.0
0.5
0.0
d2 d5 d8 d12 d2 d5 d8 d12 d2 d5 d8 d12
Zbtb10 sgRNA
Zbtb11 sgRNA
Zfp131 sgRNA
C
Zbtb10
BTB Domain
Zf C2H2 Domain
sgRNA
PAM
Endogenous
ATTCTTTGTGATGTCAGCATTGTGGTG
Ile Leu Cys Asp Val Ser Ile Val Val
Exogenous
ATTCTTTGTGATGTCTCAATCGTTGTG
ATCCTGGTTGCAGGCAGCCGTTTCTTT
Ile Leu Val Ala Gly Ser Arg Phe Phe
ATATTGGTAGCTGGAAGCCGTTTCTTT
Zbtb11
AAAGCGCACAAGTCTGTCCTCTCGGCT
Lys Ala His Lys Ser Val Leu Ser Ala
AAAGCGCACAAGAGCGTATTAAGTGCT
AAAGGAGCAGTTTCTAGTCACGAGGCT
Lys Gly Ala Val Ser Ser His Glu Ala
AAAGGAGCAGTTAGCTCCCATGAAGCT
Zfp131
CGCCACTTAATTGAGTTCACATACACA
Arg His Leu Ile Glu Phe Thr Tyr Thr
CGACATCTCATAGAATTCACATACACA
GAAGAAGAAGCCAATGATGTGTGGAAA
Glu Glu Glu Ala Asn Asp Val Trp Lys
GAGGAGGAGGCAAACGACGTATGGAAA
D
No Dox
Dox
iZbtb10-HA Zbtb10 +/+
iZbtb11-HA Zbtb11 +/+
iZfp131-HA Zfp131 +/+
E
iZbtb10-HA no dox
iZbtb10-HA dox
iZbtb11-HA dox
iZfp131-HA dox
DAPI
OCT4
HA

**Figure S5. Generation of conditional *Zbtb10Δ*, *Zbtb11Δ,* and *Zfp131Δ* lines**

**(A)** Schematic overview of the TF candidate qPCR competition assay. Non-targeting and *Zbtb10*, *Zbtb11*, and *Zfp131* targeting sgRNAs infected cells representation over 12 days of culture.

**(B)** Normalized abundance of targeting sgRNAs compared to non-targeting sgRNA measured by qPCR in the competition experiment (n=2). *Zbtb10*, *Zbtb11*, and *Zfp131* targeting sgRNAs deplete compared to non-targeting one recapitulating the TF screen depletion dynamics.

**(C)** *Zbtb10*, *Zbtb11* and *Zfp131* genes illustrated with coding regions for interdomain linker regions (gray), BTB domain (orange) and C2H2 Zinc Finger domain (green). Two sgRNAs (blue sequences) targeting PAM sites (red sequences) at the endogenous BTB domain. HA-tagged exogenous copy of ZBTB10, ZBTB11, and ZFP131 expressed at the HPRT site contain silent mutations that disable the sgRNA binding and Cas9 cutting.

**(D)** Inducible *Zbtb10-HA*, *Zbtb11-HA* and *Zfp131-HA* lines with 48 h expression of exogenous alleles. Inducing transgenes do not cause any distinct morphological changes in ESCs. Scale bar = 200 µm.

**(E)** Oct4 and HA staining of inducible *Zbtb10-HA*, *Zbtb11-HA* and *Zfp131-HA* lines with 48h expression of exogenous alleles. Inducing transgenes do not alter *Oct4* expression in ESCs. Scale bar = 100 µm.

### Figure S6

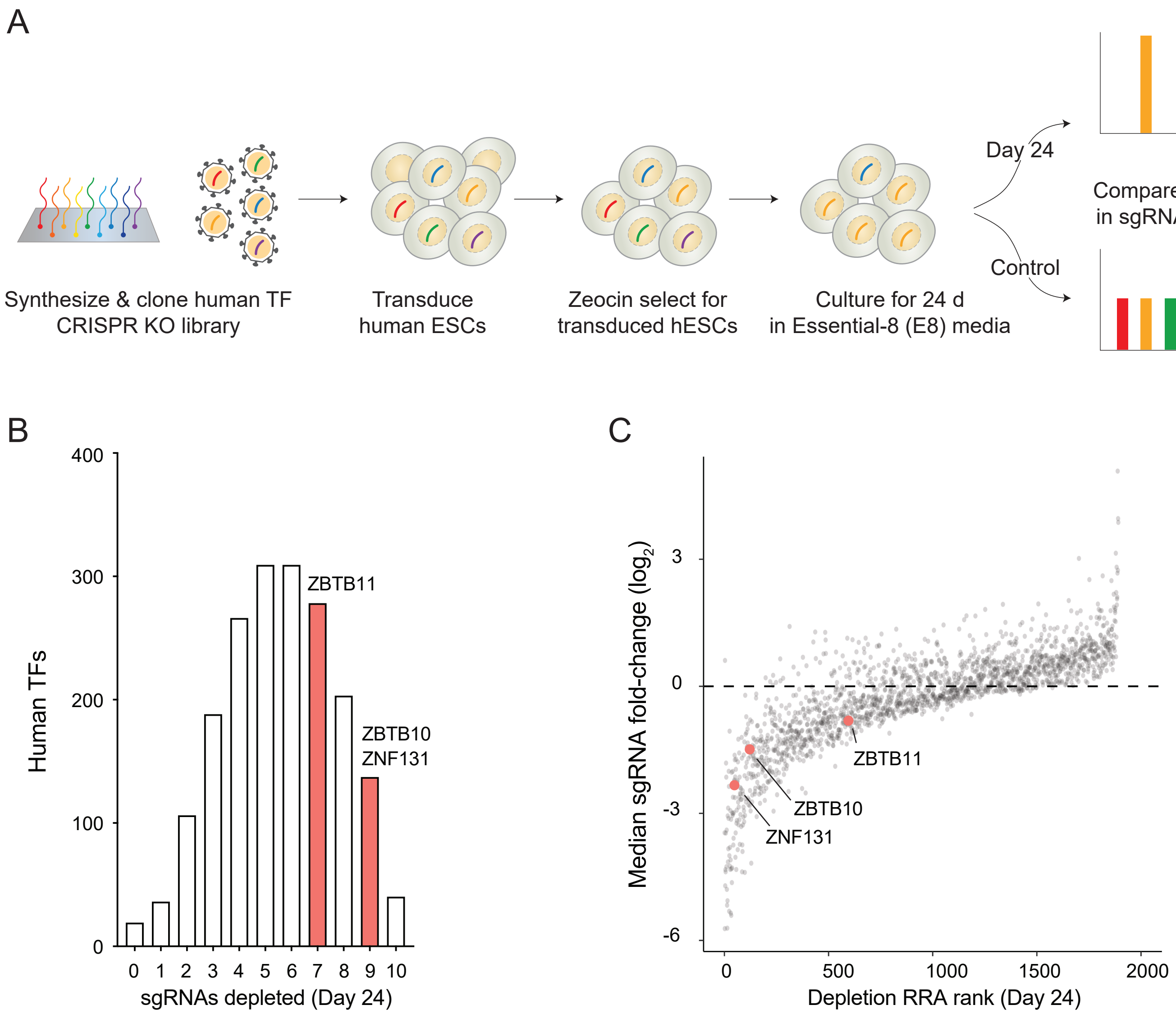
A
Day 24
Compare change
in sgRNA counts
Control
Synthesize & clone human TF
CRISPR KO library
Transduce
human ESCs
Zeocin select for
transduced hESCs
Culture for 24 d
in Essential-8 (E8) media
B
Human TFs
400
300
200
100
0
0 1 2 3 4 5 6 7 8 9 10
sgRNAs depleted (Day 24)
ZBTB11
ZBTB10
ZNF131
C
Median sgRNA fold-change (log2)
3
0
-3
-6
0 500 1000 1500 2000
Depletion RRA rank (Day 24)
ZBTB11
ZBTB10
ZNF131

**Figure S6. Human embryonic stem cell CRISPR screen identifies ZBTB10, ZBTB11 and ZNF131 as depleting genes.**

**(A)** Schematic of human embryonic stem cell screen for TFs.

**(B)** Number of depleted sgRNAs for each TF using FDR < 0.05 threshold, as calculated from the non-targeting sgRNAs. ZBTB11 has 7 depleted sgRNAs, whereas ZBTB10 and ZNF131 have 9 depleted sgRNAs in hESCs.

**(C)** Median sgRNAs fold-change (log2) at day 24 as a function of RRA gene rank. sgRNAs targeting ZBTB10, ZBTB11 and ZFP131 have lower median rank denoting their role in hESCs.

### Figure S7

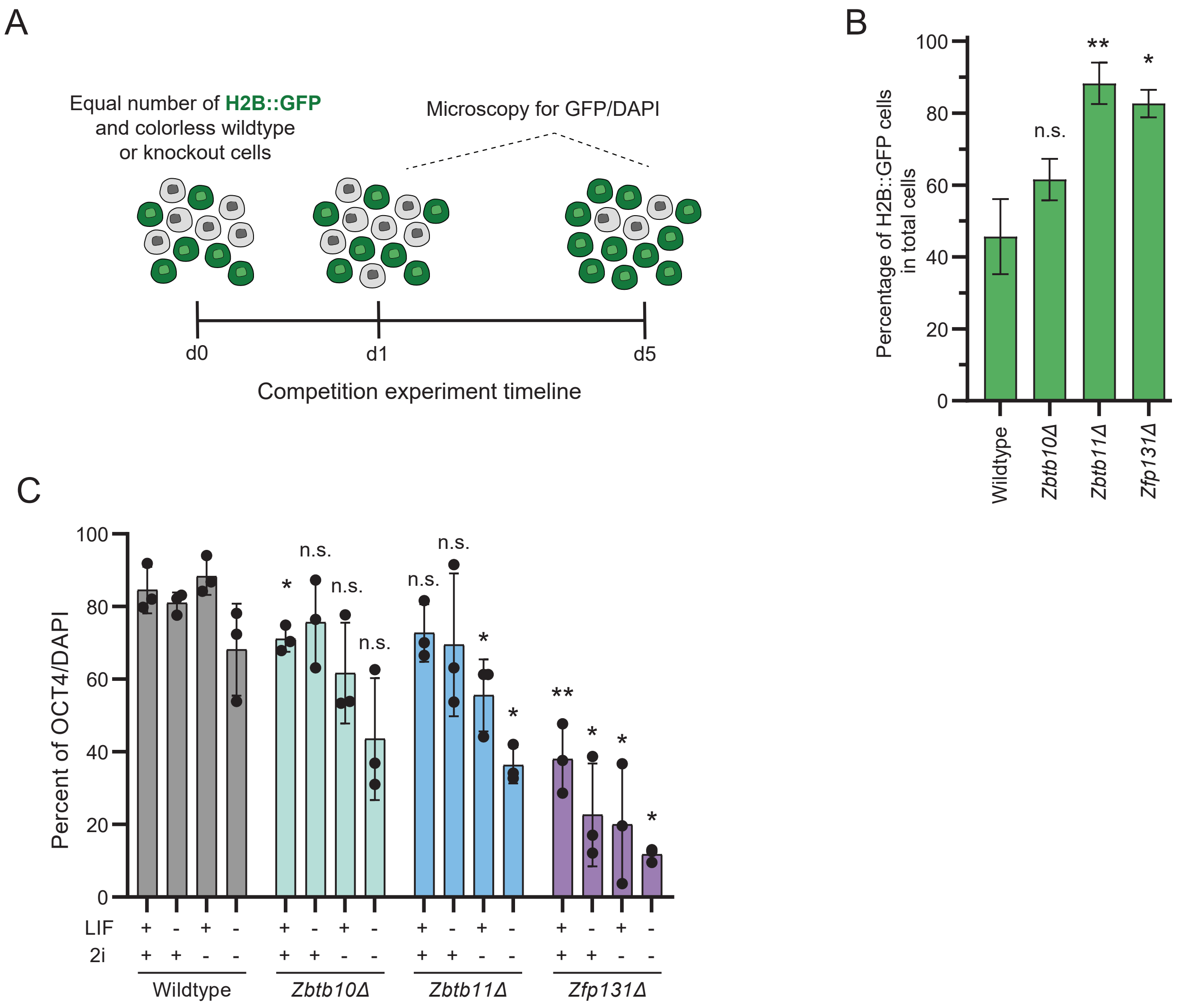
A
Equal number of H2B::GFP and colorless wildtype or knockout cells
Microscopy for GFP/DAPI
d0
d1
d5
Competition experiment timeline
B
Percentage of H2B::GFP cells in total cells
100
80
60
40
20
0
n.s.
**
*
Wildtype
Zbtb10Δ
Zbtb11Δ
Zfp131Δ
C
Percent of OCT4/DAPI
100
80
60
40
20
0
n.s.
*
**
LIF
2i
+
-
Wildtype
Zbtb10Δ
Zbtb11Δ
Zfp131Δ

**Figure S7. *Zbtb11Δ* and *Zfp131Δ* mutant cells have reduced proliferation and differentiation phenotypes.**

**(A)** Schematic overview of the cell-growth competition assay. Monoclonal mutant *Zbtb10Δ*, *Zbtb11Δ,* and *Zfp131Δ* cells compete with GFP-labeled wt ESCs.

**(B)** Percentage of non-GFP cells over time in cell growth competition assay of wt, *Zbtb10Δ*, *Zbtb11Δ,* and *Zfp131Δ* cells versus *H2b*::*Gfp* (n=3). Mirroring pooled mutant analysis, wt ESCs readily outcompete *Zbtb11Δ* and *Zfp131Δ* cells.

**(C)** OCT4/DAPI staining for wildtype, *Zbtb10Δ*, *Zbtb11Δ* and *Zfp131Δ* cells in 2i and LIF combination conditions (n=3). Statistical analysis was performed by student's t-test, *p<0.05, **p<0.005. *Zbtb11Δ* and *Zfp131Δ* cells have decreased OCT4 signal in specific culture conditions.

### Figure S8

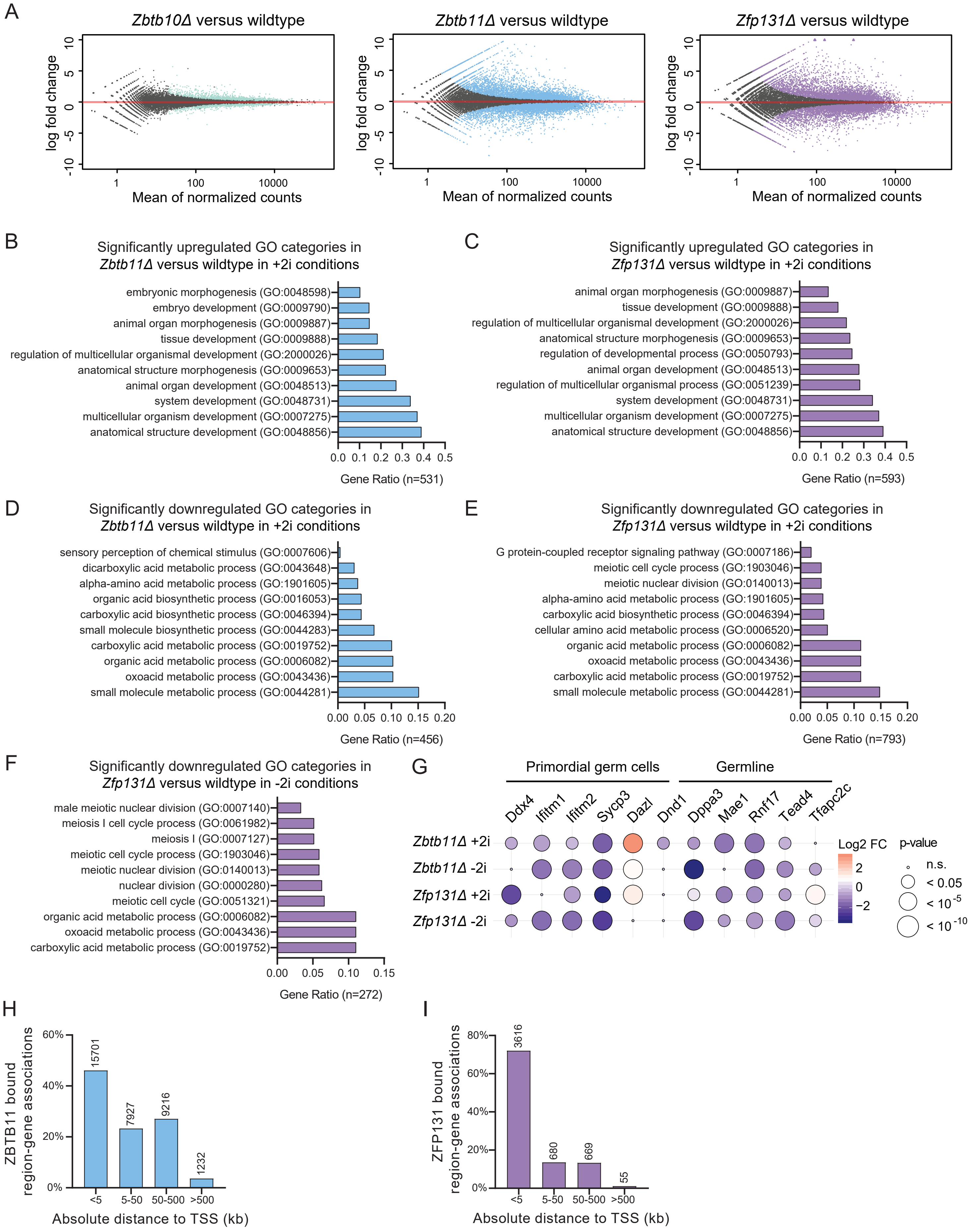

A
Zbtb10Δ versus wildtype
Zbtb11Δ versus wildtype
Zfp131Δ versus wildtype
log fold change
Mean of normalized counts
B
Significantly upregulated GO categories in Zbtb11Δ versus wildtype in +2i conditions
embryonic morphogenesis (GO:0048598)
embryo development (GO:0009790)
animal organ morphogenesis (GO:0009887)
tissue development (GO:0009888)
regulation of multicellular organismal development (GO:2000026)
anatomical structure morphogenesis (GO:0009653)
animal organ development (GO:0048513)
system development (GO:0048731)
multicellular organism development (GO:0007275)
anatomical structure development (GO:0048856)
Gene Ratio (n=531)
C
Significantly upregulated GO categories in Zfp131Δ versus wildtype in +2i conditions
animal organ morphogenesis (GO:0009887)
tissue development (GO:0009888)
regulation of multicellular organismal development (GO:2000026)
anatomical structure morphogenesis (GO:0009653)
regulation of developmental process (GO:0050793)
animal organ development (GO:0048513)
regulation of multicellular organismal process (GO:0051239)
system development (GO:0048731)
multicellular organism development (GO:0007275)
anatomical structure development (GO:0048856)
Gene Ratio (n=593)
D
Significantly downregulated GO categories in Zbtb11Δ versus wildtype in +2i conditions
sensory perception of chemical stimulus (GO:0007606)
dicarboxylic acid metabolic process (GO:0043648)
alpha-amino acid metabolic process (GO:1901605)
organic acid biosynthetic process (GO:0016053)
carboxylic acid biosynthetic process (GO:0046394)
small molecule biosynthetic process (GO:0044283)
carboxylic acid metabolic process (GO:0019752)
organic acid metabolic process (GO:0006082)
oxoacid metabolic process (GO:0043436)
small molecule metabolic process (GO:0044281)
Gene Ratio (n=456)
E
Significantly downregulated GO categories in Zfp131Δ versus wildtype in +2i conditions
G protein-coupled receptor signaling pathway (GO:0007186)
meiotic cell cycle process (GO:1903046)
meiotic nuclear division (GO:0140013)
alpha-amino acid metabolic process (GO:1901605)
carboxylic acid biosynthetic process (GO:0046394)
cellular amino acid metabolic process (GO:0006520)
organic acid metabolic process (GO:0006082)
oxoacid metabolic process (GO:0043436)
carboxylic acid metabolic process (GO:0019752)
small molecule metabolic process (GO:0044281)
Gene Ratio (n=793)
F
Significantly downregulated GO categories in Zfp131Δ versus wildtype in -2i conditions
male meiotic nuclear division (GO:0007140)
meiosis I cell cycle process (GO:0061982)
meiosis I (GO:0007127)
meiotic cell cycle process (GO:1903046)
meiotic nuclear division (GO:0140013)
nuclear division (GO:0000280)
meiotic cell cycle (GO:0051321)
organic acid metabolic process (GO:0006082)
oxoacid metabolic process (GO:0043436)
carboxylic acid metabolic process (GO:0019752)
Gene Ratio (n=272)
G
Primordial germ cells
Germline
Ddx4
Ifitm1
Ifitm2
Sycp3
Dazl
Dnd1
Dppa3
Mae1
Rnf17
Tead4
Tfapc2c
Zbtb11Δ +2i
Zbtb11Δ -2i
Zfp131Δ +2i
Zfp131Δ -2i
Log2 FC
p-value
n.s.
< 0.05
< 10-5
< 10-10
H
ZBTB11 bound region-gene associations
15701
7927
9216
1232
<5
5-50
50-500
>500
Absolute distance to TSS (kb)
I
ZFP131 bound region-gene associations
3616
680
669
55
<5
5-50
50-500
>500
Absolute distance to TSS (kb)

**Figure S8. *Zbtb11Δ* and *Zfp131Δ* mutant cells express multiple lineage-specific markers in +2i conditions.**

**(A)** Volcano plots of log2 fold change of transcripts expressed in *Zbtb10Δ*, *Zbtb11Δ* and *Zfp131Δ* cells versus wt cells grown in the absence of doxycycline for 3d in +2i+LIF media (n=2). Significant changes (* $p<0.05$) marked in color for each genotype. *Zbtb11* and *Zfp131* mutations induce aberrant gene expression.

**(B)** GO-enrichment analysis of significantly upregulated genes in *Zbtb11Δ* in +2i condition (n=531), **(C)** significantly upregulated genes in *Zfp131Δ* in +2i condition (n=593), **(D)** significantly downregulated genes in *Zbtb11Δ* in +2i condition (n=456), **(E)** significantly downregulated genes in *Zfp131Δ* in +2i condition (n=793), and **(F)** significantly downregulated genes in *Zfp131Δ* in -2i condition (n=272). Gene Ratio is calculated with the number of occurrences in specific GO categories divided by the total amount of genes. The low number of downregulated genes in -2i conditions for *Zbtb11Δ* produced no significant GO-term enriched.

**(G)** The log2 fold change of TFs obtained from bulk RNAseq experiment in *Zbtb11Δ* and *Zfp131Δ* cells versus wildtype in plus and minus 2i conditions. (n.s. Not significant, * $< 0.05$, ** $< 10^{-5}$, *** $< 10^{-10}$) (n=2). No aberrant expression of primordial germ cells and germline TFs detected in *Zbtb11Δ* and *Zfp131Δ* cells.

**(H)** The proximity of all ZBTB11 and **(I)** ZFP131 bound regions to TSS. Both TFs bind proximal to TSSs.

### Figure S9

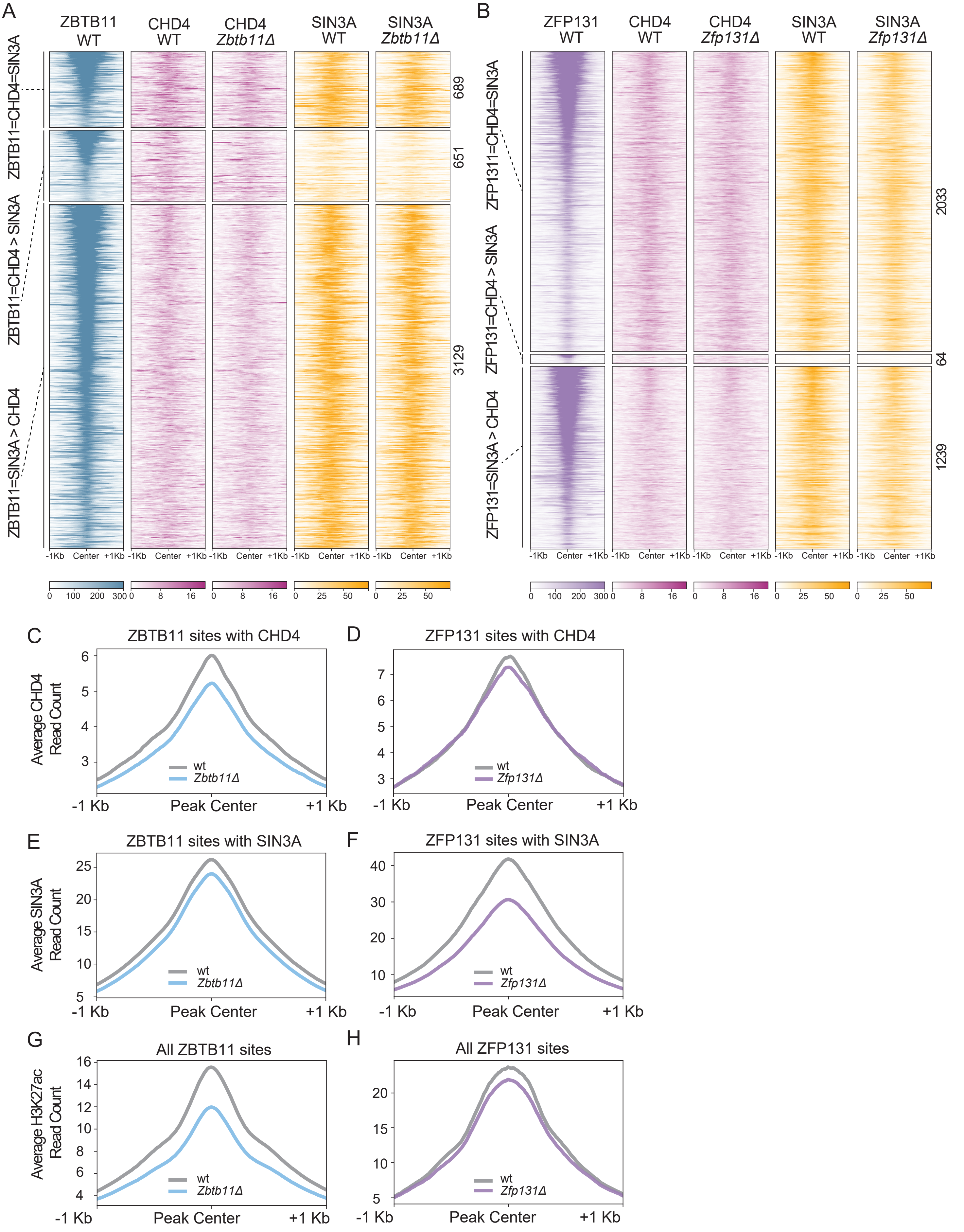

A
ZBTB11 WT
CHD4 WT
CHD4 *Zbtb11Δ*
SIN3A WT
SIN3A *Zbtb11Δ*
ZBTB11=CHD4=SIN3A
ZBTB11=CHD4 > SIN3A
ZBTB11=SIN3A > CHD4
689
651
3129
-1Kb Center +1Kb
0 100 200 300
0 8 16
0 25 50
B
ZFP131 WT
CHD4 WT
CHD4 *Zfp131Δ*
SIN3A WT
SIN3A *Zfp131Δ*
ZFP1311=CHD4=SIN3A
ZFP131=CHD4 > SIN3A
ZFP131=SIN3A > CHD4
2033
64
1239
C
ZBTB11 sites with CHD4
Average CHD4 Read Count
wt
*Zbtb11Δ*
-1 Kb
Peak Center
+1 Kb
D
ZFP131 sites with CHD4
*Zfp131Δ*
E
ZBTB11 sites with SIN3A
Average SIN3A Read Count
F
ZFP131 sites with SIN3A
G
All ZBTB11 sites
Average H3K27ac Read Count
H
All ZFP131 sites

**Figure S9. ZBTB11 and ZFP131 cobound with CHD4 and SIN3A.**

**(A)** ChIP-seq heatmap for ZBTB11 overlap with CHD4 and SIN3A in *Zbtb11Δ* and wt cells (n=2 for all).

**(B)** ChIP-seq heatmap for ZFP131 overlap with CHD4 and SIN3A in *Zfp131Δ* and wt cells (n=2 for all).

**(C)** Metagene plots for CHD4 binding difference in *Zbtb11Δ* versus wt (n=2) and in **(D)** *Zfp131Δ* versus wt (n=2). Both mutants have a slight decrease in CHD4 binding in mutant vs wt.

**(E)** Metagene plots for SIN3A binding difference in *Zbtb11Δ* versus wt (n=2) and in **(F)** *Zfp131Δ* versus wt (n=2). Both mutants have a slight decrease in SIN3A binding in mutant vs wt.

**(G)** Metagene plots for H3K27ac at ZBTB11 bound sites in *Zbtb11Δ* and wt ESCs (n=2) and **(H)** at ZFP131 bound sites in *Zfp131Δ* versus wt (n=2). Both mutants have a decrease in H3K27ac contrary to the expected increase.
